# DeepVir: A reproducible workflow for large-scale viral dark matter discovery

**DOI:** 10.64898/2026.09.15.751837

**Authors:** Matheus Augusto Calvano Cosentino, Nicolas Fernandez, Marcelo A. Soares, Ahidjo Ayouba, André Felipe Andrade dos Santos, Mirela D’arc

## Abstract

High-throughput sequencing (HTS) has revolutionized virosphere exploration. However, characterizing highly divergent viral sequences remains a bottleneck known as Viral Dark Matter (VDM). Numerous tools were developed to unravel such diversity, but they usually require complex prior HTS data analysis processes. Consequently, a common bottleneck to VDM exploration is the manual, chained execution of complex command-line applications. To address the need for automated and scalable viral discovery, we developed DeepVir, a reproducible Snakemake pipeline that integrates classical homology-based alignments with profile Hidden Markov Model (HMM) mining of the RNA-dependent RNA polymerase (RdRp). To validate the pipeline’s efficacy, we analyzed 385.23 GB of publicly available transcriptomic data (203 Sequence Read Archive libraries) from 49 American bat species. DeepVir successfully identified 179 distinct viral groups. This included 903 contigs spanning nine known viral families, enabling the characterization of novel genomes within *Orthomyxoviridae* (Influenza A H7N9), *Picornaviridae*, *Alphaflexiviridae*, *Retroviridae* (*Spumaretrovirinae*), *Papillomaviridae*, *Herpesviridae*, and *Adenoviridae*. Furthermore, the pipeline uncovered 170 putative novel VDM lineages. By employing deep homology searches and Sequence Similarity Network (SSN) visualization, we contextualized these highly divergent VDM sequences, revealing significant evolutionary relationships with the orders *Mononegavirales* and *Bunyavirales*. Notably, human-driven curation of the pipeline’s outputs allowed for the cross-library assembly of the first putative exogenous *Spumavirus* in the Americas. Ultimately, by automating complex bioinformatic processing steps, scalable pipelines like DeepVir empower researchers to prioritize the biological and epidemiological interpretation of their findings, accelerating the characterization of wildlife virospheres and enhancing pathogen genomic surveillance.

## 1. Introduction

With the development and global implementation of high-throughput sequencing (HTS) protocols, our understanding of viral diversity has improved significantly, revealing viruses as the most abundant and diverse entities in the biosphere ^1^. As of January 1st, 1994, at the onset of HTS development ^2^, 15,375 viral sequences were available in the public GenBank database from the National Center for Biotechnology Information (NCBI), rising to 13,018,162 by January 1, 2024, an 846-fold increase in 30 years. Beyond curated assemblies, this same period highlights an unprecedented availability of raw genomic data: the Sequence Read Archive (SRA) hosts over 29.5 million sequencing runs, comprising approximately 89 petabases of sequencing data. Consequently, this abundance of data allowed us to unravel the complex ecological dynamics between viruses and their hosts. Beyond their role as etiological agents ^3,4^, viruses also contribute to immune system maturation ^5^ and play key roles in biogeochemical events ^6^. However, it has also exposed a novel bottleneck in viral identification known as Viral Dark Matter (VDM), comprising sequences generated by HTS that remain unidentifiable due to the high genetic divergence from known viruses in genetic databases ^7,8^.

The high evolutionary rate of RNA viruses ^9,10^, combined with the absence of universal marker genes, complicates the identification of divergent viruses present within the HTS data. Current strategies increasingly rely on protein structure and Hidden Markov Model (HMM) profiles rather than local sequence alignments, which are up to 10 times more conserved ^11^. Nowadays, the RNA-dependent RNA polymerase (RdRp) is the closest approximation to a universal marker within *Riboviria* ^12^, serving as a key reference for phylogeny and modern classification of these RNA viruses that lack a DNA stage ^7,12,13^. Recent initiatives ^7,13–15,16^ have demonstrated the immense potential of viral discovery within the SRA by mining over 10 petabases of public sequencing data and discovering 10^5^ novel RNA viruses, expanding the number of known species by nearly an order of magnitude. By analyzing millions of sequencing libraries using highly optimized alignment and HMM architectures, these massive-scale efforts have uncovered a hidden RNA virosphere ^7,13–15,16^. Despite those findings, the applicability of VDM identification methodologies for non-bioinformatics experts remains a hurdle. State-of-the-art tools and pipelines designed to explore the VDM often depend on specialized cloud computing infrastructures, which are often impractical for everyday laboratory routines. Alternatively, when available for local use, they require complex manual HTS data analysis prior to the execution of the tool of interest ^7,13,14,17^. Consequently, a major bottleneck preventing a global approach to viral discovery is the manual, chained execution of complex command-line applications, which alienates researchers lacking advanced computational expertise.

Workflow engines like Snakemake address the abovementioned challenge by automating pipelines and ensuring reproducibility ^17^. This effectively democratizes access to sophisticated genomic analysis, empowering researchers across diverse regions to contribute to genomic studies. In this work, we present DeepVir, a custom Snakemake pipeline designed to integrate HTS classical viral homology search complemented with HMM profile RdRp mining. DeepVir is a Snakemake workflow designed to democratize and automate the initial steps of viral discovery. It targets virologists seeking a reproducible, scalable, and fault-tolerant pipeline to work within viral metagenomics. By leveraging Snakemake, DeepVir provides efficient task parallelization to avoid queue bottlenecks, suspend-and-resume capabilities for handling job failures, and comprehensive data logging ^17^. This allows researchers to focus on biological interpretation rather than pipeline development and management.

To validate the pipeline scalability, we reanalyzed publicly available transcriptome and meta-transcriptome datasets of American bats from the NCBI SRA. Previous investigations on free living world populations identified diverse viral groups naturally infecting bats, notably Ebola-related viruses ^18–21^, coronaviruses ^22–24^, and arboviruses such as Dengue and Zika viruses ^25,26^. The use of HTS ^27,28^ and mining of publicly available HTS data ^13,29^ has revealed that the currently known bat virus diversity represents only a fraction of the broader virosphere ^30–32^. Furthermore, its systematic application in American bat surveillance remains limited as HTS studies to explore their Virosphere are scarce, relying mostly on PCR methodologies to detect viral agents with a limited genomic representation. By mining American bat host-derived transcriptomic data, we aimed to uncover the VDM of the American bats to enhance global surveillance of reservoir species harboring potentially high-risk zoonotic viruses.

## 2. Material and Methods

### 2.1. DeepVir: Viral Discovery Reproducible Pipeline

For viral discovery in the HTS data of interest, DeepVir was developed as an automated and modular Snakemake (v.9.11.1) pipeline with strict dependency isolation across environments, via Conda channel management ^33^. Each analytical tool is associated with a dedicated YAML environment specification file, ensuring that all software dependencies, versions, and libraries are strictly encapsulated and portable across different platforms. To scale efficiently within High-Performance Computing (HPC) infrastructures, DeepVir natively integrates the snakemake-executor-plugin-slurm v.2.^34^, submitting each individual rule as an isolated cluster job with optimized resource allocation. The core architecture of DeepVir is compartmentalized into four distinct operational modules and a centralized reporting layer, which can be dynamically triggered by command-line flags depending on the experimental scope **(Figure 1)**.

**Figure 1.**
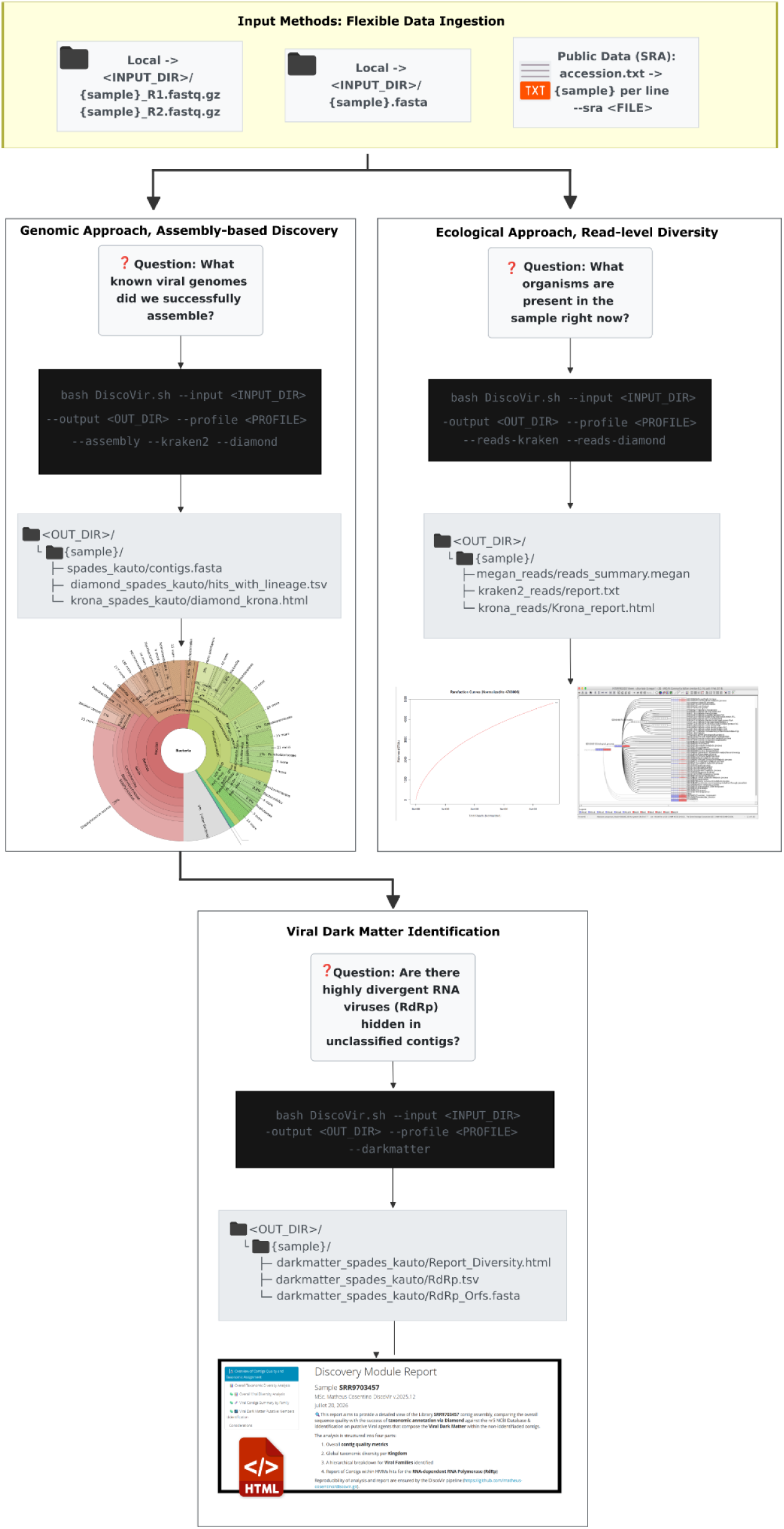
Conceptual architecture of the DeepVir pipeline. The workflow is driven by specific biological questions and organized into modular analytical tracks. The Input Phase supports flexible data ingestion, accepting local paired-end reads, pre-assembled contigs, or automated fetching of public data via NCBI SRA accessions. The Ecological Approach module (activated via --reads-kraken and --reads-diamond) performs read-level taxonomic profiling to quickly assess overall sample composition, generating MEGAN summaries and rarefaction curves. The Assembly-Based Diversity module (--assembly) conducts *de novo* assembly and assigns taxonomy to the resulting contigs, outputting standardized TSV lineage tables and interactive Krona plots. The Viral Dark Matter Exploration module (--darkmatter) uses the unclassified contigs generated in the assembly step to mine for highly divergent RNA viruses by identifying RdRp domains, summarizing the findings in an interactive RMarkdown HTML report. Black terminal boxes illustrate the exact bash execution commands required to trigger each module, while grey directory boxes display the resulting reproducible output folder structure.

#### Raw Data & Pre-processing

The pipeline accepts local raw sequencing data (paired-end or single-end FASTQ formats within the directory given via the --input flag) or automates public data retrieval. By providing a line-separated text file containing Sequence Read Archive (SRA) identifiers via the --sra flag, DeepVir executes downloads using sra-tools v.3.1.0 ^35^. Ingested raw reads are subsequently subjected to adapter trimming and quality and length filtering (>Q30 and >50 nt, respectively) utilizing Fastp v.0.20.1 ^36^ to yield clean data for downstream exploration.

#### Ecological Approach, Read-level Diversity

Designed for rapid pathogen identification and community ecology profiling directly from read-level data, this module bypasses assembly to query filtered reads. Taxonomic profiling can be executed using Kraken2 v.2.1.3 ^37^ (activated via --reads_kraken) and Diamond v.2.1.13 ^38^ (activated via --reads_diamond). Read-level taxonomy generated by Diamond is meganized to enable downstream inspection using MEGAN v.6.25.10 ^39^. Abundance tables are automatically formatted into BIOM files, and interactive hierarchical community structures are generated using Krona v.2.7.1 ^40^, alongside a rarefaction curve estimation to evaluate sequencing depth.

#### Genomic Approach, Assembly-based Discovery

To enable the recovery of partial or near-complete viral genomes from complex biological data, high-quality filtered reads are directed to the *de novo* assembly stage using the --assembly flag. To accommodate both metagenomic and metatranscriptomic studies, DeepVir allows selection among assemblers. For short-read datasets, users can select either Megahit v.1.2.9 ^41^ or SPAdes v.3.15.3^42^, both of which are easily configured within the pipeline. DeepVir allows dynamic selection of full suite of SPAdes algorithmic modes (including meta-, metaviral-, and rnaviralSPAdes) and the targeted k-mer size strategies (either automated or user-defined array inputs) to optimize graph construction based on the viral target type.

Following assembly, the reconstructed contigs undergo a taxonomic profiling workflow highly symmetrical to the read-level analysis to characterize the assembled community. Assembled sequences can be rapidly profiled using Kraken2 v.2.1.3 (activated by the --kraken2 flag), which yields interactive hierarchical plots generated by Krona v.2.7.1. As a complementary option, contigs are characterized through Diamond v.2.1.13 (activated by the --diamond flag) via local alignments against the protein database. The resulting hits pass through a taxonomic refinement step using TaxonKit v.0.20.0 ^43^ to append official lineages derived from the NCBI taxonomy database. Finally, these comprehensive outputs are translated into interactive Krona diversity plots and meganized file formats to enable comparative downstream exploration in MEGAN v.6.25.10 ^39^.

#### Viral Dark Matter Identification

Assembled contigs without a taxonomic assignment in Diamond are further explored to identify putative RNA viruses within the VDM in HTS data (activated by the --darkmatter flag). Orphan contigs have their Open Reading Frames (ORFs) translated into amino acid sequences using getorf Emboss v.6.6.0 ^44^, with a minimum length cut-off of 200 amino acids applied across all viral genetic codes (translation tables 1, 3, 4, 5, 6, 11, and 16)^7^. Redundant sequences are removed using CD-HIT v.4.8.1^45^ with a 90% identity cut off. The remaining non-redundant amino acid sequences are scanned for canonical RdRp palm domains via HMMER v.3.3.2 ^46^ utilizing palm_annot scripts ^47^.

**Table 1.**
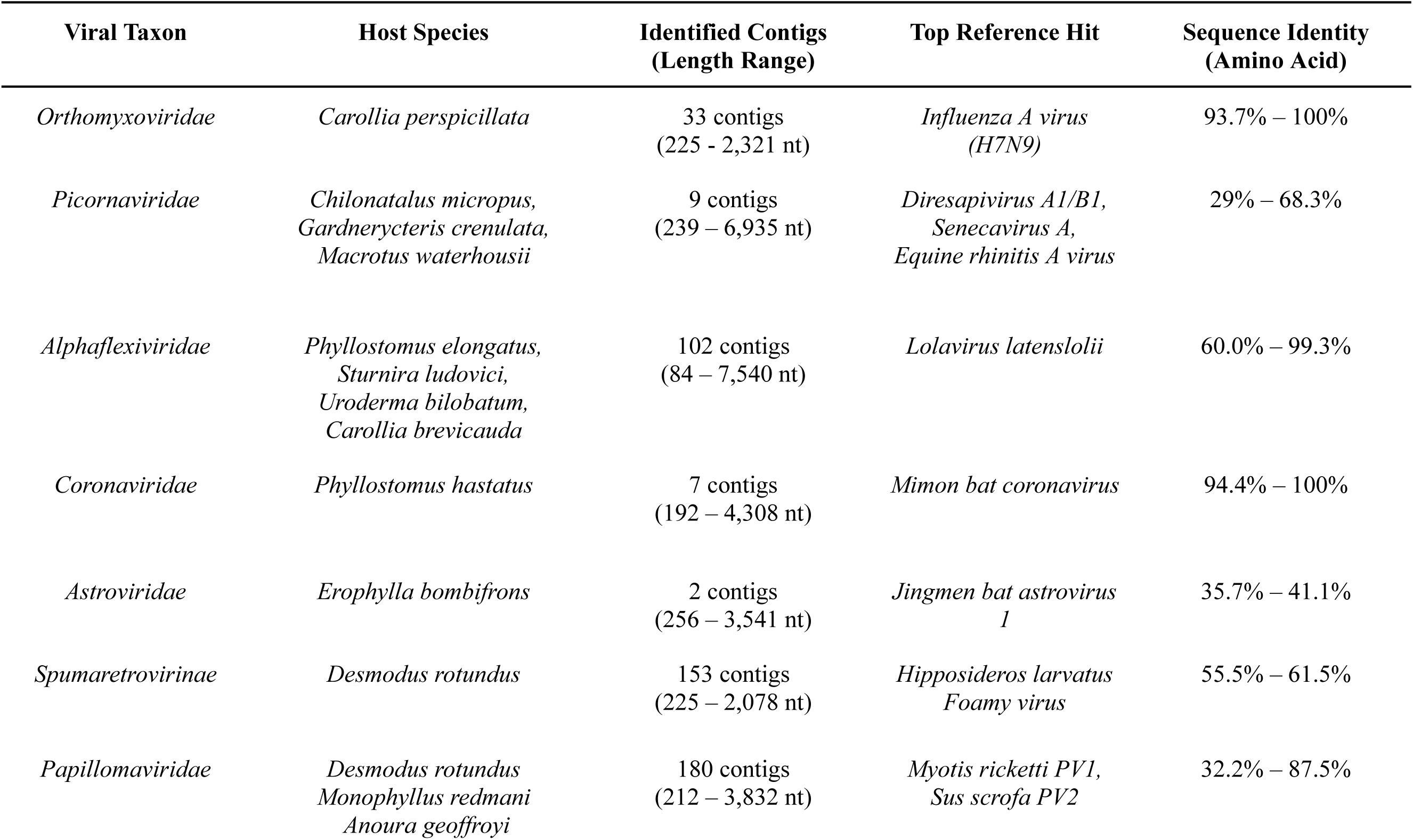
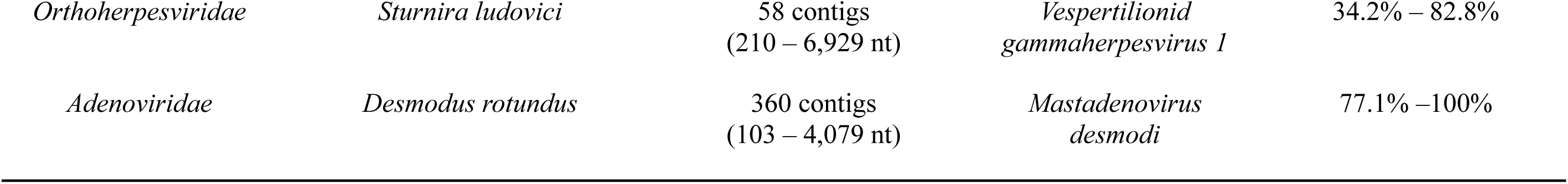
Summary of novel viruses identified via the DeepVir pipeline by Homology based search (DIAMOND BlastX).

#### Reporting

Outputs across all active modules converge into unified dashboards **(Figure 2)**. Pre-processing metrics are aggregated via MultiQC v.1.32 ^48^, taxonomic diversity is explored through interactive hierarchical Krona plots, and final viral diversity profiles including putative VDM candidates are summarized into programmatically compiled HTML reports synthesized via R v.4.5.2 and the rmarkdown v.2.30 package.

**Figure 2.**
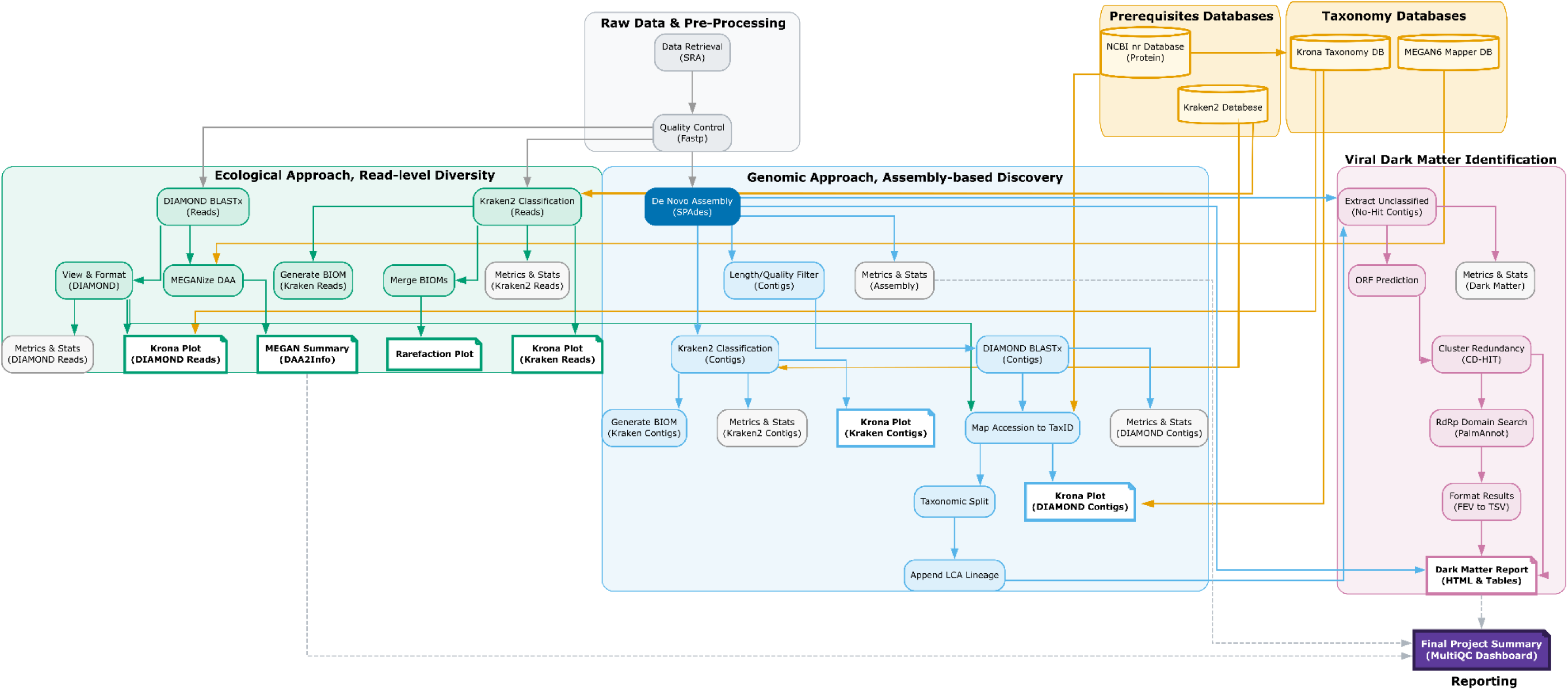
Schematic directed acyclic graph of the DeepVir pipeline for viral discovery. The pipeline architecture is organized into distinct, interconnected functional modules: (0) Prerequisites & Taxonomy Databases (Yellow), representing the external prerequisite reference databases (NCBI nr, Kraken2) and taxonomy databases download by the pipeline on the first run (Krona taxonomy, and MEGAN mapping) required for downstream assignments; (1) Raw Data & Pre-Processing (Grey), where raw sequencing data is retrieved from the NCBI SRA and filtered for quality using Fastp; (2) Ecological Approach, Read-level Diversity (Green), which performs rapid read-level taxonomic profiling via Kraken2 and Diamond BLASTx, generating MEGAN-based summaries, BIOM tables, and rarefaction plots; (3) Genomic Approach, Assembly-based Discovery (Blue), responsible for de novo assembly (e.g., SPAdes), contig length/quality filtering, and robust lineage refinement using custom mapping to append Lowest Common Ancestor (LCA) taxonomic lineages; and (4) Viral Dark Matter Identification (Pink), which mines taxonomically unclassified (no-hit) contigs for RdRp palm domains using ORF prediction, sequence redundancy clustering (CD-HIT), and HMM validation (palm_annot). Intermediate metrics and statistics from all active modules ultimately converge into a Final Project Summary (Purple) unified MultiQC dashboard. Cylindrical icons indicate reference databases, standard rectangular boxes indicate processing rules, and note-shaped icons denote primary tabular and interactive reporting outputs (e.g., Krona plots, HTML reports). Solid arrows represent standard data flow, while yellow lines trace the incorporation of reference database information into analytical steps.

#### Prerequisites & Taxonomy Databases

For database management, the reference databases for Kraken2 and Diamond must be pre-installed by the user. However, the taxonomic databases required for Krona, MEGAN, and TaxonKit are automatically downloaded and configured during the pipeline’s first run. This automated process requires internet connection and approximately 50 GB of available local storage (∼29 GB for Krona, ∼9.5 GB for MEGAN, and ∼12 GB for TaxonKit/NCBI taxonomy files).

### 2.2. Biological Validation of DeepVir

To evaluate the performance and scalability of the DeepVir pipeline, we selected paired-end RNA-sequencing data from free-living American bats from meta-transcriptomics and transcriptomic BioProjects available at SRA (https://www.ncbi.nlm.nih.gov/sra), featuring a median of ∼10 million reads per library **(Supplementary Table S1)**. To investigate the presence of putative novel or highly divergent viruses previously unreported in the original studies, the workflow was run using the --assembly flag configured to use the rnaviralSPAdes v.3.15.3 ^49^ algorithm, which removes low-complexity reads typical of RNA assemblies during graph construction. Concurrently, the --diamond flag was deployed to scan assembled contigs for shared homologies against the complete NCBI non-redundant (nr) protein database v.5. (https://ftp.ncbi.nlm.nih.gov/blast/db/v5/). Finally, the --darkmatter module was activated to screen unassigned orphan contigs using palm_annot scripts to pinpoint highly divergent viral RdRp signatures that presented a minimum RdRp score of 50. All viral candidates identified through this multi-layered pipeline underwent rigorous biological validation, including structural homology assessments, confirmatory read back-mapping, and maximum-likelihood phylogenetic analyses to establish their evolutionary context, as detailed in **Supporting Information Appendix S1.**

A total of 385.23 GB of raw sequencing data was processed, comprising 203 paired-end SRA datasets from 49 species and seven families of the Chiroptera order (**Supplementary Table S1**). The dataset was dominated by the family Phyllostomidae (n = 171), with *Desmodus rotundus* being the most represented species (n = 72; 35.5%) **(Supporting Information Appendix S2: Supplementary Figure S1**). While hematophagy was the most represented dietary niche due to its sole representative, *D. rotundus*, the dataset showed greater species diversity among frugivores (n = 59, across 20 species) and insectivores (n = 43, across 17 species). Sample sources were heterogeneous, encompassing feces, saliva, and vomeronasal epithelium, whereas heart, eye, and olfactory epithelium tissues were the most common across all taxa.

## 3. Results

### 3.1. Viral Diversity

By applying the DeepVir pipeline into the public HTS data, our analysis identified 179 distinct viral groups, which were categorized into two main classes: those found by homology-based local alignments (n = 9; 5.02%) and those found by HMM RdRp search (n = 170; 94.98%) **(Figure 3A**, **Table 1)**. The known diversity comprised 903 viral contigs with clear homology to nine previously reported viral families. In contrast, 182 contigs (representing 170 putative novel groups) were classified as VDM due to the lack of a recognizable taxonomic assignment **(Supplementary Table S2)**. For the known viral contigs, the identification of major marker genes allowed for direct phylogenetic characterization (detailed in subsequent sections).

**Figure 3.**
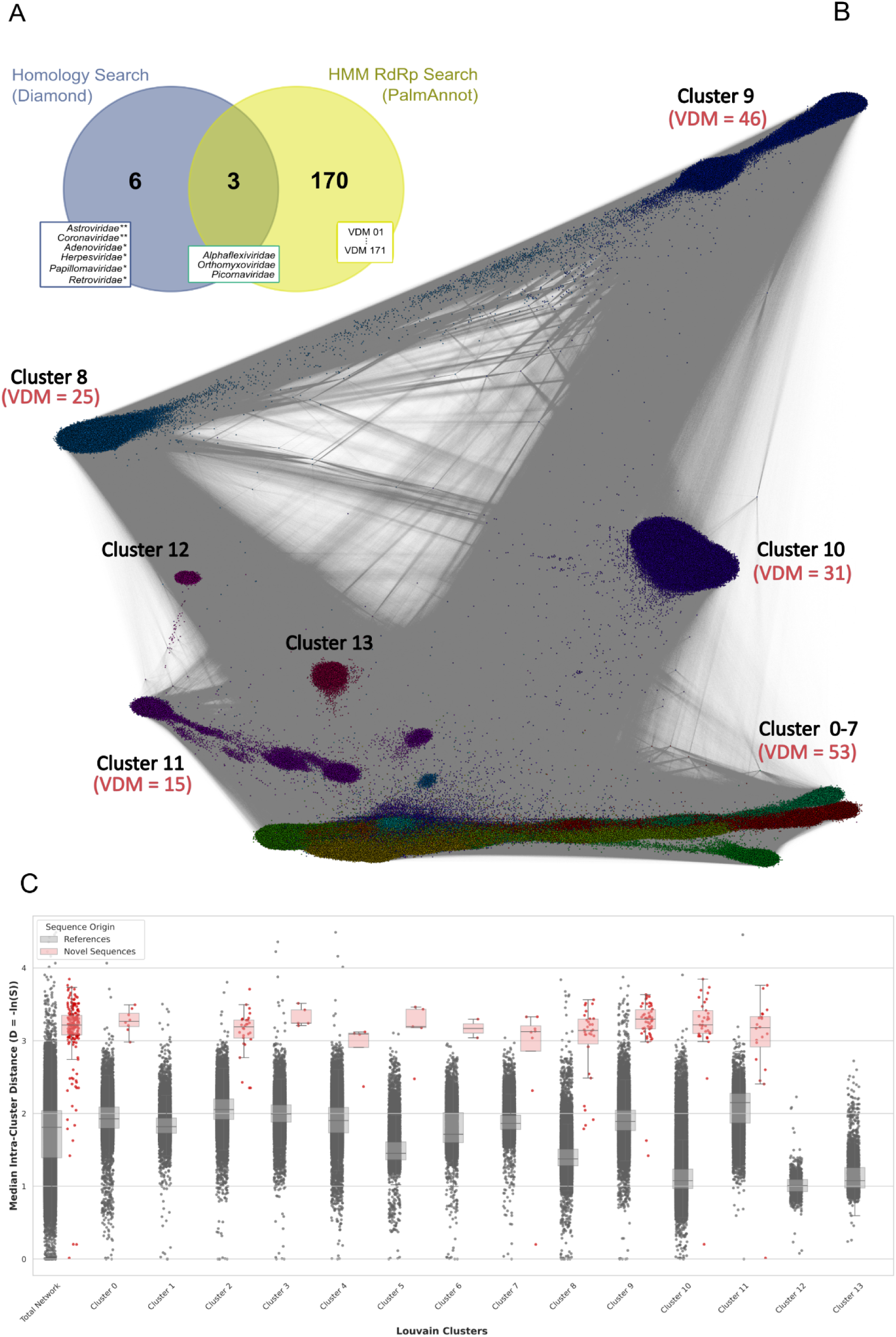
Summarized viral diversity identified and evolutionary divergence of novel VDM sequences. (A) Venn diagram illustrating all identified viral contigs per homology search strategy implemented. Viral groups marked (*) do not belong to the Riboviria realm, while marked (**) are Riboviria detected despite the absence of RdRp protein in HTS data. (B) Sequence Similarity Network (SSN) based on structural homology. Nodes represent individual viral sequences, grouped into 14 highly supported Louvain clusters. (C) Boxplots and overlaid dot plots comparing the median intra-cluster distances of known viral references from public databases (grey boxes) and the newly identified viral dark matter contigs (red points) across the same clusters. In panel C, boxplots indicate the median (central line) and the interquartile range, while individual sequences are overlaid as jittered points to illustrate data density and highlight divergent outliers.

Conversely, to contextualize the highly divergent VDM contigs within known viral diversity, we applied a two-tiered classification strategy. First, a palmID-based search identified only 11 VDM contigs sharing significant homology with PalmDB representatives **(Supporting Information Appendix S2: Figure S2 & S3)**. The low success rate of this approach (11/182; 6.04%), despite the confirmed presence of RdRp domains by the Palm_annot scripts **(Supplementary Table S2)**, underscores the extreme evolutionary divergence of these sequences from established viral clades. To overcome this limitation, we employed an MMseqs2 deep homology search coupled with SSN visualization. This approach successfully assigned all RdRp palm domains (513,368 nodes connected by 86,860,058 edges) into 14 highly supported Louvain clusters (modularity = 0.6760) **(Figure 3B)**. These novel VDM members demonstrated a substantial genetic distance from known viral diversity in public databases (min: 0.758, max: 2.7679) **(Figure 3C)**. By evaluating the dominant annotated taxonomic orders and families within each cluster, we were able to infer putative host associations and ecological niches for these novel viruses **(Supporting Information Appendix S2: Supplementary Figures S4 & S5, Supplementary Table S2)**.

Notably, the most present contigs were found in fungal-associated clusters (Clusters 8 and 9, with 30 and 50 contigs, respectively), where *Cryppavirales*, *Ourlivirales*, and *Wolframvirales* were the dominant taxa. Altogether, bacteriophage-associated sequences were also abundant, exemplified by Cluster 10, which consisted of 33 contigs nested within a large-scale cluster of over 29,000 *Levivirales* sequences **(Supplementary Table S2)**. Two clusters (6 and 12) remained unassigned, reflecting the presence of highly divergent viral lineages yet to be explored. In the vertebrate-vector interface, Cluster 11 was particularly relevant, comprising 20 contigs alongside a high prevalence of *Mononegavirales* (1,775 sequences) and *Bunyavirales* (1,454 sequences) **(Supplementary Table S2)**.

### 3.2. Characterization of Assembled Viral Genomes

#### Orthomyxoviridae

*De novo* assembly of metagenomic data from libraries SRR10059480 and SRR19391903, which originated from the same individual of *Carollia perspicillata,* (Biosample SAMN12675088, BioProject PRJNA563501), identified 33 *Orthomyxoviridae* contigs. Homology searches confirmed these as distinct genomic segments encoding the polymerase complex (PB2, PB1, PA), hemagglutinin (HA), neuraminidase (NA), nucleocapsid (NP), and matrix proteins, with sequence identity ranging from 93.7% to 100% **(Supplementary Table S3)**. Among contigs, three were identified as PB1. Phylogenetic analysis placed them within the diversity of Influenza A virus (as members of genus *Alphainfluenzavirus*) with strong support (SH-aLRT = 100 & UFBoot = 100) **(Supporting Information Appendix S2: Supplementary Figure S6)**, while HA and NA genotyping confirmed the subtype as H7N9 **(Supporting Information Appendix S2: Supplementary Figure S7)**. Reference-based assembly revealed a low number of reads mapped (395 for SRR10059480 and 242 for SRR19391903). However, the horizontal genomic coverage was high for both, 92.6% and 82.1%, respectively.

#### Picornaviridae

A total of nine assembled contigs across three libraries were identified as belonging to the ssRNA+ family *Picornaviridae*, with assembly ranging from 239 to 6,935 nt. Among these, a near-complete genome (6,935 nt) was identified in an eye transcriptomics library of *Chilonatalus micropus* (SRR9703457, Bioproject PRJNA555243) **(Figure 4A)**, containing complete coding regions for the P1 and 3Dpol proteins, which are key to characterization of this putative novel virus. Furthermore, partial sequences showing significant homology to P1 and 3Dpol proteins were identified in another eye transcriptome library from *Macrotus waterhousii* (SRR9703483, BioProject PRJNA555243) and in an olfactory epithelium transcriptome from *Gardnerycteris crenulata* (SRR19391887, BioProject PRJNA563501) **(Supplementary Table S3)**. The identified regions were successfully aligned against ICTV *Picornaviridae* P1 and 3Dpol reference alignments to characterize these putative novel viruses within the diversity of the family *Picornaviridae*. By gene prediction and homology sharing identification, a total of three libraries containing five contigs classified as putative novel *Picornaviridae* were used for phylogenetic inference, which successfully recreated known genus with strong support (SH-aLRT and UFBoot > 75). No clear host virus co-divergence topology was identified as no homology by the host group was visualized **(Figure 4B & 4C)**.

**Figure 4.**
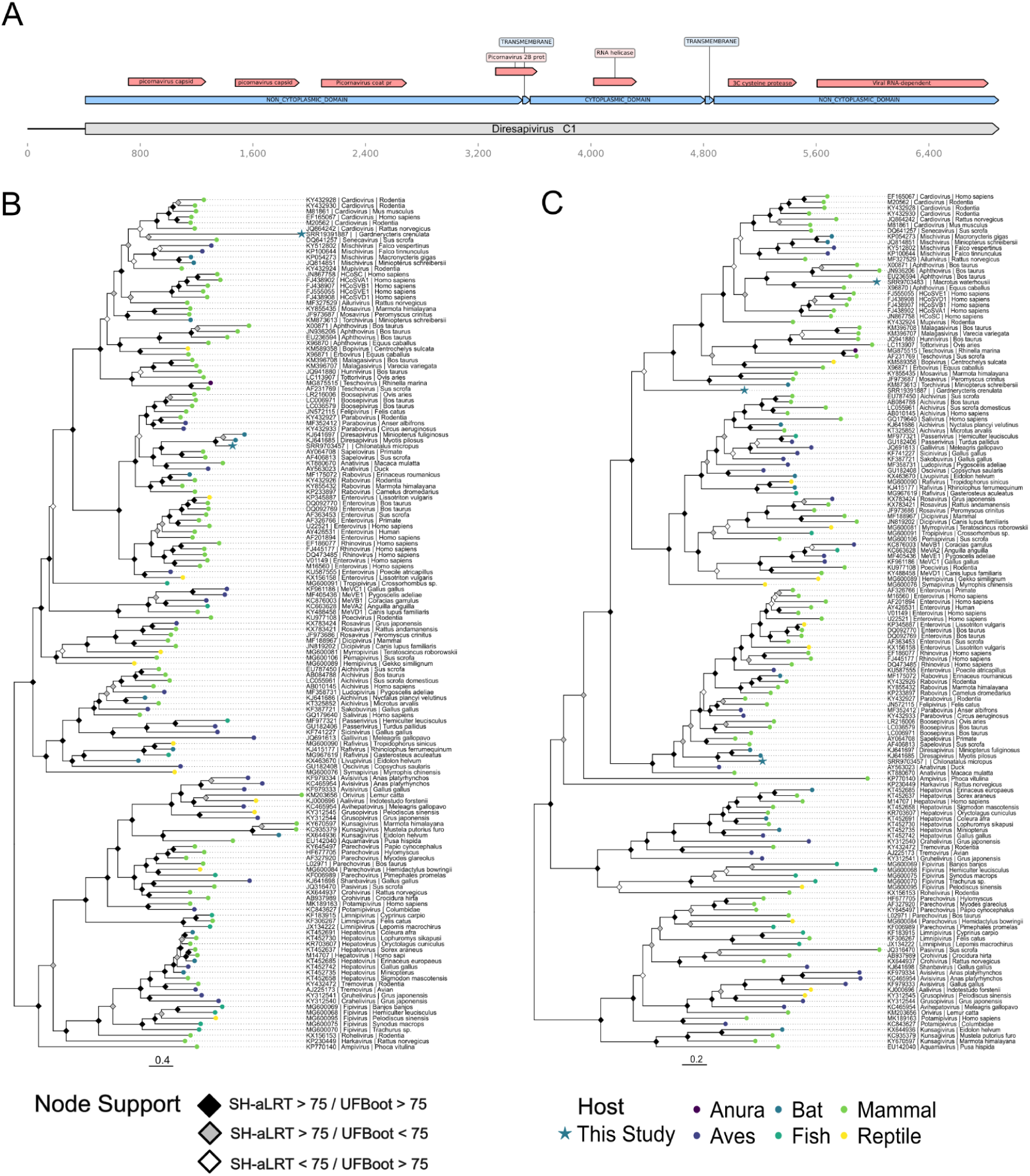
Novel picornaviruses identified in American bats. (A) Schematic representation of Diresapivirus C1 genome. The diagram displays the predicted Open Reading Frame (ORFs; positions 411–6,899) encoding a single polyprotein. Colored annotations indicate conserved domains identified via Pfam/InterPro, including: the structural capsid module (rhv-like/P1) at the N-terminus, followed by the non-structural domains Helicase (Superfamily 3), 3C Cysteine Protease, and the RNA-dependent RNA polymerase (RdRp/3Dpol) at the C-terminus. The scale bar represents nucleotide length. (B-C) Maximum likelihood phylogenetic tree of the family Picornaviridae, based on the P1 capsid protein (B) and the 3Dpol RNA-dependent RNA polymerase (C). Phylogenies were inferred from an alignment of 149 sequences (1,267 amino acid sites) using the Q.PFAM+F+I+R8 model for P1, and 150 sequences (533 amino acid sites) using the Q.PFAM+F+I+R7 model for 3Dpol. Tips are colored according to the host group: Anura (dark purple), Bats (blue), Aves (dark teal), Fish (dark green), Mammals (light green), and Reptiles (yellow). Sequences generated in this study are marked with stars. Node support is indicated by diamonds: black diamonds represent high support (SH-aLRT > 75 and UFBoot > 75), gray diamonds represent support > 75 in SH-aLRT only, and white diamonds in UFBoot only. The scale bar indicates the number of amino acid substitutions per site.

The near-complete genome identified in *Chilonatalus micropus* (SRR9703457, BioProject PRJNA555243) clustered with high support (3Dpol & P1, SH-aLRT = 100 and UFBoot = 100) as a sister lineage to the cluster formed by Chinese bat picornaviruses, *Diresapivirus A1* (KJ641685) and *B1* (KJ641697). This novel virus shares amino acid identities of 64.5–68.3% (P1) and 64.7–65.6% (3Dpol) with those viruses (**Supplementary Table S4**) and has been provisionally named *Diresapivirus C1*, representing the putative third species within this genus.

Regarding the *Picornaviridae* contigs found in the olfactory epithelium transcriptome of *Gardnerycteris crenulata* (SRR19391887, BioProject PRJNA563501), partial fragments of P1 (214 aa, 642 nt) and 3Dpol (468 aa, 1,404 nt) were recovered. Phylogenetic reconstruction retrieved the 3Dpol fragment as a divergent lineage with moderate support (SH-aLRT = 59.2; UFBoot = 84), positioned outside the major clades comprising genera such as *Cardiovirus*, *Aphthovirus*, *Teschovirus*, and *Mosavirus*. This distinct position is supported by the low amino acid identity (33.7–39.8%) shared with members of these genera (**Supplementary Table S4**). Phylogenetic reconstruction of the P1 gene retrieved the sequence as a sister group to *Senecavirus A* with moderate support (SH-aLRT = 85.1; UFBoot = 66), forming a clade distinct from, but related to the genus *Cardiovirus* **(Figure 4C)**. Despite this clustering, the sequence shares only 30.9% amino acid identity with *Senecavirus A*.

Finally, among the *Picornaviridae* contigs identified in *Macrotus waterhousii* (SRR9703483), only a partial fragment corresponding to 3Dpol was recovered (278 aa, 834 nt). Phylogenetic reconstruction placed the sequence within the genus *Aphthovirus*, clustering as a sister group to *Equine rhinitis A virus* (ERAV) with high support (SH-aLRT = 92.6; UFBoot = 99) and sharing 53.7% amino acid identity **(Figure 4C & Supplementary Table S4)**.

#### Alphaflexiviridae

A total of 102 contigs belonging to the family *Alphaflexiviridae* were identified across four SRA libraries. These contigs, ranging in length from 84 to 7,540 nt, were detected in four bat species sampled in Peru under BioProject PRJNA563501, which investigated chemosensory evolution in bats. The host species were *Phyllostomus elongatus*, *Sturnira ludovici*, *Uroderma bilobatum*, and *Carollia brevicauda*. Diamond BLASTx analysis revealed that all contigs shared moderate-to-high identity (60–99.3%) with *Lolavirus latenslolii* (EU489641) **(Supplementary Table S3)**. Furthermore, eight contigs containing both replicase and capsid domains, which are essential for viral classification were selected for contextualization. Phylogenetic analysis placed these contigs within the *Lolavirus latenslolii* clade (EU489641) with high branch support for both proteins (SH-aLRT = 100; UFBoot = 100) **(Figure 5)**. These contigs showed high amino acid identity with *L. latenslolii* for the replicase (97.7–98.6%) and capsid (87.5–93.5%) domains, while the novel strains demonstrated intra-group variation of 96.6–100% for the replicase and 79.9–100% for the capsid **(Supplementary Table S4)**.

**Figure 5.**
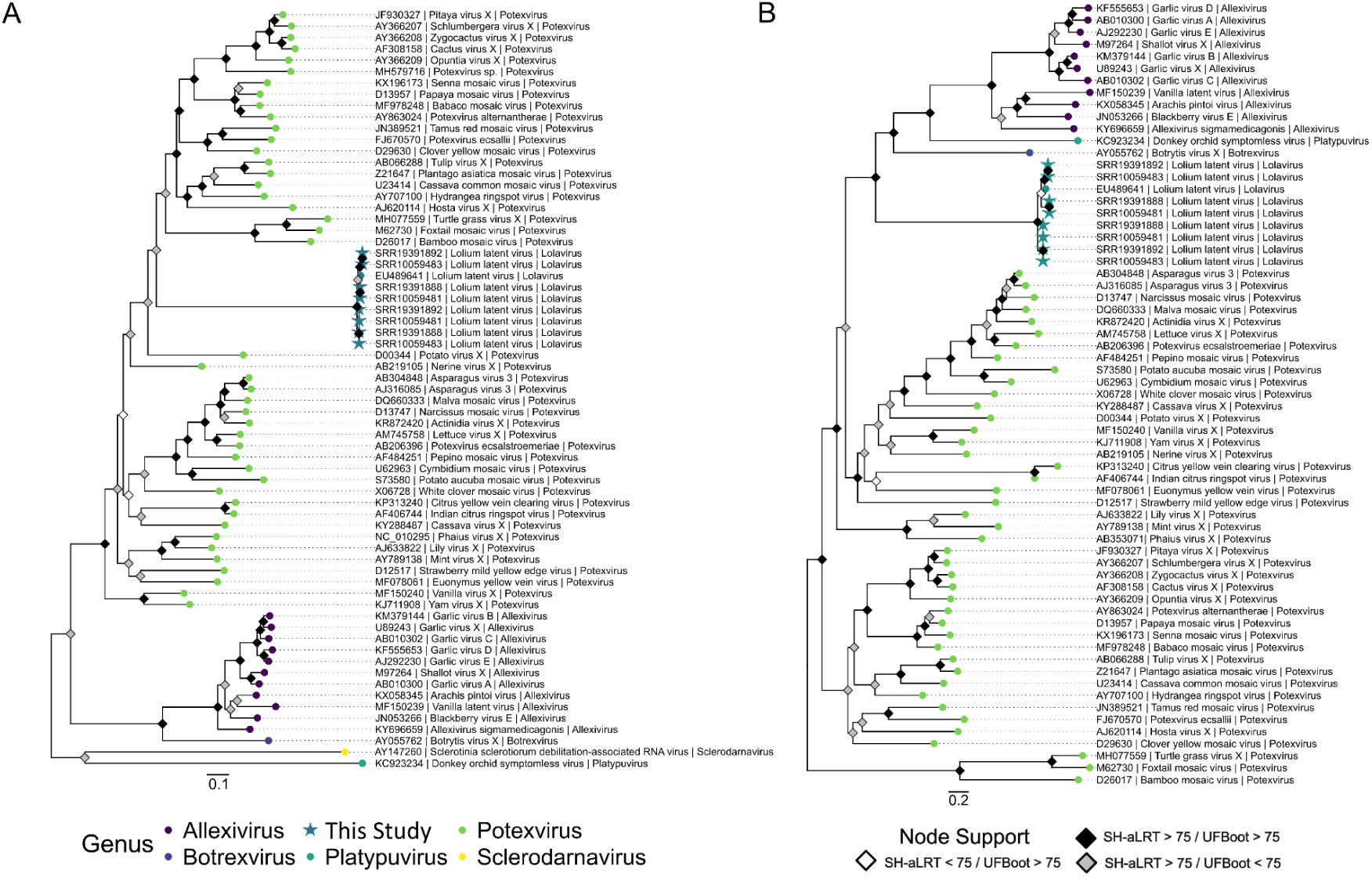
Novel reports of *Lolavirus* identified in American bats. (A-B) Maximum likelihood phylogenetic tree of the family Alphaflexiviridae, based on the Replicase protein (A) and the Capsid (B). Phylogenies were inferred from an alignment of 67 sequences (878 amino acid sites) using the LG+F+R6 model for Replicase protein, and 65 sequences (140 amino acid sites) using the Q.PFAM+G4 model for Capsid. Tips are colored according to the current ICTV viral genus: *Allexivirus* (dark purple), *Botrexvirus* (blue), *Lolavirus* (dark teal), *Platypuvirus* (dark green), *Potexvirus* (light green), and *Sclerodarnavirus* (yellow). Sequences generated in this study are marked with stars. Node support is indicated by diamonds: black diamonds represent high support (SH-aLRT > 75 and UFBoot > 75), gray diamonds represent support > 75 in SH-aLRT only, and white diamonds in UFBoot only. The scale bar indicates the number of amino acid substitutions per site.

#### Coronaviridae

Among the assembled contigs, seven were identified as belonging to the *Coronaviridae* in a *Phyllostomus hastatus* adult male eye RNA-seq library (SRR9703480, BioProject PRJNA555243). A disparity in their quality and size was observed, as one contig stands out with a length of 4,308 nt and high vertical coverage (31.4x), whereas the other six contigs were shorter (ranging from 192 to 648 nt) and exhibited very low vertical coverage (from 1.25x to 2.56x). Nevertheless, BLASTn analysis identified homology of the contigs with high nucleotide identity (87.37–94.66%) within the spike, nucleocapsid, NS3, and matrix genes of the previously described *Mimon bat coronavirus isolate PREDICT/PDF-3316* (MZ293744) **(Supporting Information Appendix S2: Supplementary Figure S8)**.

#### Astroviridae

Within the assembled contigs, two were identified as belonging to the family *Astroviridae* in an *Erophylla bombifrons* eye RNA-seq library (SRR9703488, BioProject PRJNA555243). A disparity in their quality and size was observed, as one contig stands out with a length of 3,541 nt and a coverage of 7.88x, whereas the second contig was considerably shorter (256 nt) and exhibited very low coverage (1.43x). Furthermore, Diamond BLASTx analysis identified the contigs’ homology with a low amino acid identity (35.71– 41.18%) within the transmembrane domains of the *Jingmen bat astrovirus 1* (OQ802697) **(Supporting Information Appendix S2: Supplementary Figure S8)**.

#### Spumaretrovirinae

Among the assembled contigs, 153 were identified as belonging to the family *Retroviridae*, subfamily *Spumaretrovirinae*. Such assemblies were distributed among 35 libraries from BioProjects PRJEB28138 and PRJEB34487, studies that explored the metatranscriptomes of the common vampire bat, *D. rotundus*. Contigs’ sizes varied with a minimum length of 225 nt, and a maximum length of 2,078 nt. A total of 65 contigs were successfully mapped to a reference against the recently reported *Spumaretrovirinae* of *Hipposideros larvatus* bats (OR951385), with an assembled consensus estimated at 9,378 nt and an estimated genome coverage of 79.43% **(Figure 6A)**. Annotation transfer from Geneious identified coding sequences corresponding to *gag*, partial *RT,* and partial *env* coding sequences, while the Prokka identified two putative complete accessory genes, *tas* and *bet*.

**Figure 6.**
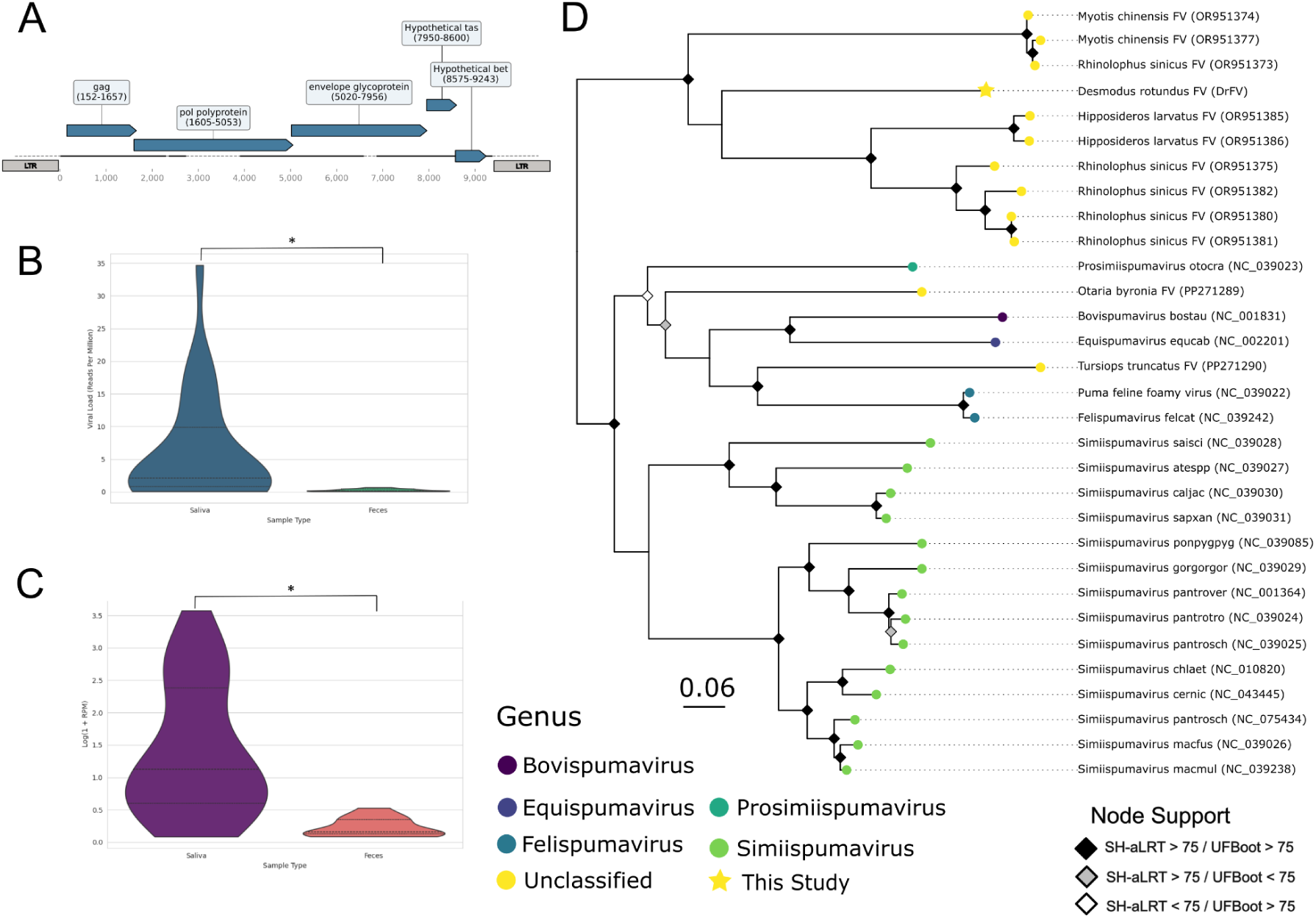
Genomic and phylogenetic characterization of the *Desmodus rotundus* Foamy virus (DrFV). (A) Genomic organization of the assembled DrFV consensus sequence. The predicted coding sequences for the Gag, Pol, and Envelope (Env) proteins are shown as blue arrows. The positions of the putative accessory genes, *tas* and *bet*, are also indicated. Regions without representation are shown as dashed lines. (B-C) Comparison of estimated viral load between saliva and feces samples. Estimated viral load is quantified as Reads Per Million (RPM) mapped to the viral consensus. The violin plots show the distribution of the normalized RPM (B) and log-transformed (C) viral load, illustrating significant higher viral loads in saliva compared to feces (*, p < 0.0001). (D) Maximum likelihood phylogenetic tree of the *Spumaretrovirinae* subfamily, illustrating the phylogenetic placement of the novel foamyvirus found in *Desmodus rotundus*. Phylogeny was inferred with a 708 amino acids alignment under the model rtREV+I+G4. The tips of the tree are color-coded to represent the different viral genera: *Bovispumavirus* (purple), *Equispumavirus* (dark blue), *Felispumavirus* (light blue), *Simiispumavirus* (green), *Prosimiispumavirus* (dark green), and unclassified genus (yellow). Node support is indicated by diamonds: black diamonds represent high support (SH-aLRT > 75 and UFBoot > 75), gray represent support in SH-aLRT only, and white in UFBoot only. The scale bar indicates the number of amino acid substitutions per site.

The consensus sequence assembled was used as a reference to map the filtered reads. Estimated viral load was significantly higher in saliva (RPM: mean = 6.00, range = 0.09 - 34.69; Log-RPM: mean = 1.41, range = 1.41 - 3.57) compared to feces (RPM: mean = 0.26, range = 0.27 - 0.70; Log-RPM: mean = 0.23, range = 0.23 - 0.53), as determined by the Mann-Whitney U test (p < 0.0001) **(Figure 6B & 6C)**. The 708-residue amino acid alignment of the polymerase protein revealed pairwise amino acid identities between previously described spumaretroviruses and *Desmodus rotundus foamy virus* (DrFV) ranging from 55.5% to 61.5% (mean: 58.15%). The phylogenetic inference recovered a highly supported monophyletic lineage of unclassified bat *Spumaretrovirinae* as a sister clade within previously reported *Spumaretrovirinae* lineages (SH-aLRT = 100, UFBoot = 100) **(Figure 6D)**. DrFV was inferred as an outgroup to previously reported *Hipposideros* and *Rhinolophus Foamyviruses* with low branch support (SH-aLRT = 25.7, UFBoot = 57), indicative of an uncertain phylogenetic reconstruction.

#### Papillomaviridae

A total of 180 assembled contigs were identified as belonging to the family *Papillomaviridae* (PV), with lengths ranging from 212 to 3,832 bp and a mean length of 562 bp. Libraries identified belonged to the BioProjects on the common vampire bat metatranscriptomics from Peru (PRJEB28138 and PRJEB34487, n = 24) and from libraries developed in comparative studies of eye transcriptomes of American bats (PRJNA555243, n = 2). Contigs were subjected to a local alignment step against custom-built databases of Diamond BLASTx for genes *E1*, *E2*, *L1* and *L2* **(Supplementary Table S3)**. Hits showed high variation in lengths, from 96 to 1,503 nt, and the sequence identity spanned from 32.2% to 87.5%. To avoid index-hopping effects within the BioProjects libraries in further characterization steps, only contigs longer than 600 bp identified as *L1* were considered, resulting in 12 libraries within informative contigs. The identified *L1* genes were phylogenetically contextualized within the current PAVE database genomes as on April 1, 2025 (https://pave.niaid.nih.gov).

The inferred *L1* tree identified PVs contigs as belonging to three distinct lineages (Supporting Information Appendix S2: **Supplementary Figure S9**). The well supported PV lineage, provisionally designated *Desmodus rotundus PV 1* (DrPV1; SH-aLRT = 99.9, UFBoot = 100) was identified in libraries ERR3569479, ERR2756795, ERR3569475, ERR3569485, ERR3569477, and ERR2756797. Sequences shared a high amino acid identity among them (95.6–100%) and DrPV1 was inferred as a sister lineage to the *Myotis ricketti PV 1* (JQ814847) with strong branch support (SH-aLRT = 89.5, UFBoot = 96) and sharing 77.27–82% amino acid identity. Together, these form a well-supported lineage (SH-aLRT = 99.8, UFBoot = 100) that is sister to the genera *Omegapillomavirus*, *Dyopapillomavirus,* and *Alphapapillomavirus*.

The second lineage, provisionally designated *Desmodus rotundus PV 2* (DrPV2; SH-aLRT = 99.9, UFBoot = 100) found within ERR3569477, ERR2756792, and ERR2756786 libraries, showed a high identity of amino acid (100%) among sequences. This lineage was inferred as a divergent lineage within the clade formed by *Rhopapillomavirus* and *Dyoomegapapillomavirus* (SH-aLRT = 80.1, UFBoot = 85). The DrPV2 showed pairwise amino acid identify values ranging from 64.9 to 76.3% relative to previously characterized *Rhopapillomavirus* and *Dyoomegapapillomavirus*, respectively.

The third identified lineage in this study was a monophyletic group formed by *Desmodus rotundus PV 3* (DrPV3; ERR2756804) and *Monophyllus redmani PV 1* (MredPV1, SRR9703484), also with strong support (SH-aLRT = 91.4, UFBoot = 76). Both DrPV3 and MredPV1 share 87.8% of pairwise amino acid identity and were inferred as a sister lineage to the *Dyophipapillomavirus Talpa europaea PV 1*, which shares 83.17% and 71.7% identity with DrPV3 and MredPV1, respectively.

Three libraries with the presence of partial genes of *L1*, *L2*, *E1* and *E2* were identified: ERR3569477 and ERR2756792 from *Desmodus rotundus*; and SRR9703484 from *Monophyllus redmani*. A concatenated phylogeny based on the identified region for the four genes was inferred, and all the near-complete viral genomes were identified as putative representatives to the three lineages **(Figure 7)**. Among the identified strains, only DrPV2 demonstrated a distinct pattern of clustering in its concatenated tree, being inferred as a sister lineage to *Sus scrofa PV 2* (SsPV2, genus *Dyothetapapillomavirus*) with strong branch support (SH-aLRT = 82.8, UFBoot = 89). The inferred genetic distance from the concatenated tree, together with the low shared amino acid identity between these viruses and their closest relatives **(Table 2)** and other previously characterized PVs **(Supplementary Table S4)**, support their classification as putative representatives of a novel genera within the family *Papillomaviridae*.

**Figure 7.**
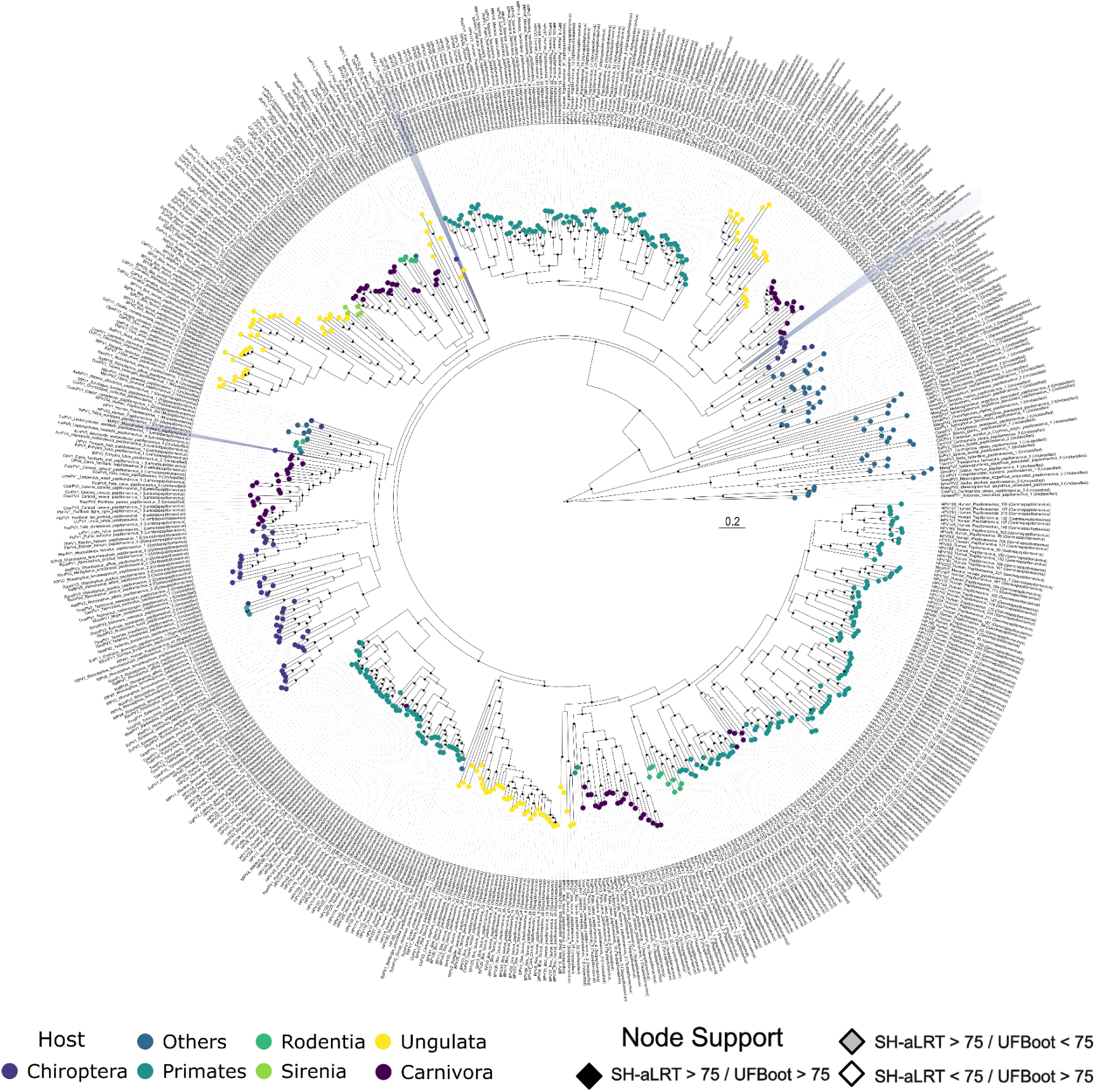
Concatenated phylogenetic tree of novel papillomaviruses found within American bats. Maximum likelihood phylogenetic tree of the Papillomaviridae family, based on concatenated L1, L2, E1 and E2 genes. Viruses found in the present work are highlighted in dark blue and stars. Phylogeny was inferred with an alignment of 602 sequences and 2092 amino acids under the model Q.yeast+F+I+G4. The tips of the tree are color-coded to represent different hosts: Chiroptera (dark blue), Carnivora (purple), Others (blue), Primates (teal), Rodentia (dark green) and Sirenia (light green). Node support is indicated by diamonds: black diamonds represent high support (SH-aLRT > 75 and UFBoot > 75), gray diamonds represent support superior to 75 in SH-aLRT only, and white diamonds in UFBoot only. The scale bar indicates the number of amino acid substitutions per site.

**Table 2.** Pairwise identity of novel PVs within closest sequences in amino acid per genes.

| Query | L1 | L2 | E1 | E2 | Closest Sequences |
| --- | --- | --- | --- | --- | --- |
| DrPV1 | 78.3% | 71.1% | 52.4% | 42.9% | MrPV1<br>(JQ814847) |
| DrPV2 | 76.3% | 58.8% | 60.4% | 43.6% | SsPV2<br>(KY817993) |
| MrPV1 | 71.7% | 44.9% | 54.6% | 47% | TePV1<br>(KC460987) |

#### Herpesviridae

A total of 58 contigs belonging to the family *Orthoherpesviridae* were identified in library SRR10059483, originating from *Sturnira ludovici* from Peru (BioProject PRJNA563501), from a vomeronasal epithelium sample. The contigs ranged in size from 210 bp to 6,929 bp, with a mean length of 1,660 bp. Among assembled contigs, 25 containing predicted genes were identified, which were concatenated to form a scaffold of 75,219 bp. Among those contigs, ten contigs containing the six core genes used in the *Orthoherpesviridae* taxonomy by ICTV were identified using local Diamond BLASTx analyses **(Supplementary Table S3)**. Maximum likelihood phylogeny of the concatenated core genes placed the novel herpesvirus within the subfamily *Gammaherpesvirinae*, as a sister lineage of the genus *Percavirus* with high support (SH-aLRT = 94.8, UFBoot = 98) **(Figure 8)**. The virus was provisionally named *Sturnira ludovici gammaherpesvirus*. This phylogenetic placement, together with the pairwise amino acid identity of the core genes **(Supplementary Table S4)** supports *Sturnira ludovici Gammaherpesvirus* as a member of a putative novel genus within the subfamily *Gammaherpesvirinae*.

**Figure 8.**
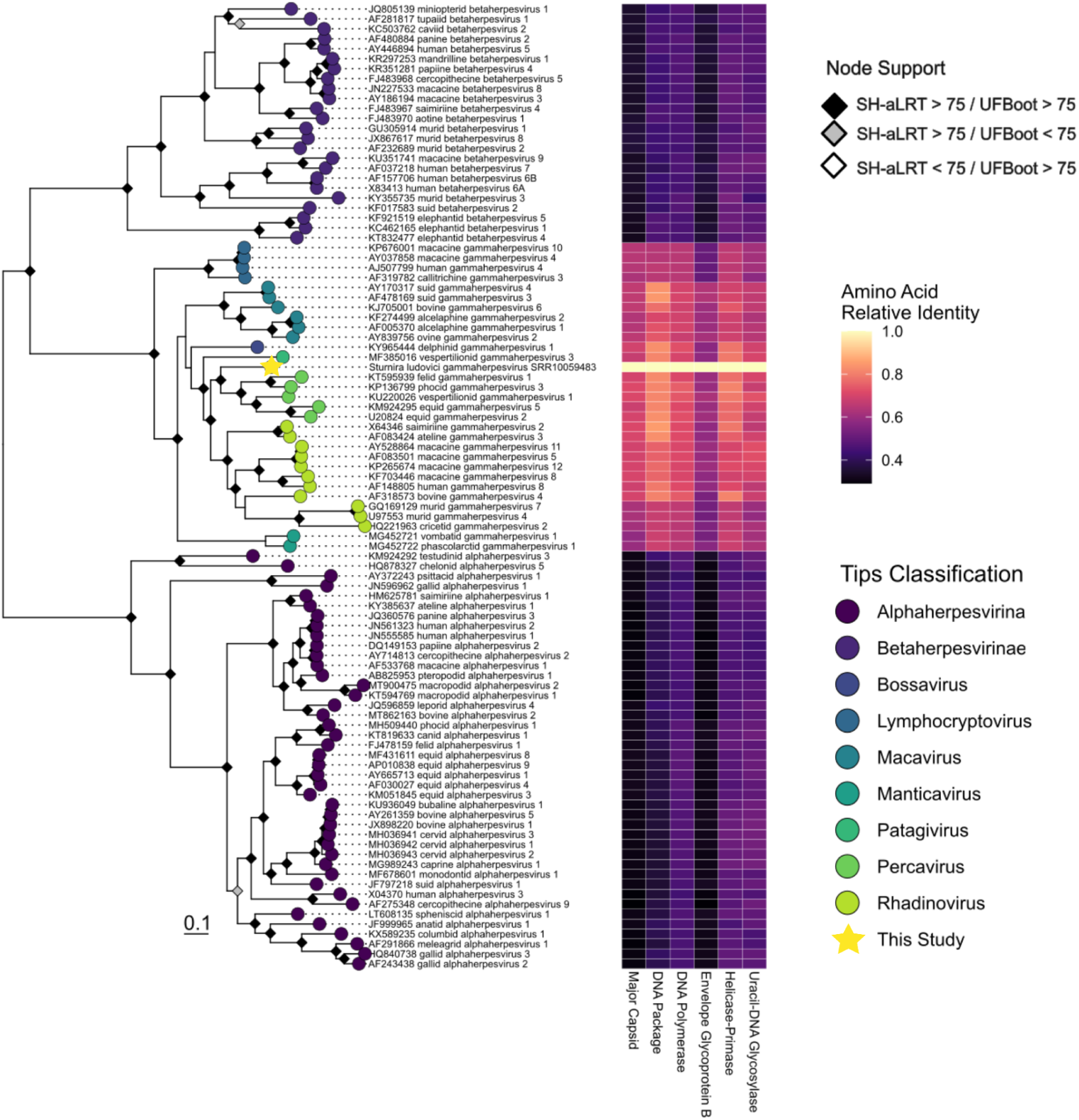
Concatenated phylogenetic tree of novel *Sturnira ludovici Gammaherpesvirina*e. A maximum likelihood phylogenetic tree of the Orthoherpesviridae family, based on the Major Capsid DNA Package, Helicase-Primase, Envelope Glycoprotein B, DNA Polymerase, Uracil-DNA Glycosylase concatenated genes. The phylogeny was inferred with an alignment of 97 sequences and 3087 amino acids under the model Q.INSECT+R7. The tips of the phylogenetic tree are classified as: Alphaherpesvirinae (dark purple), Betaherpesvirinae (blue), Bossavirus (light blue), Lymphocryptovirus (dark teal), Macavirus (green), Manticavirus (mint green), Patagivirus (sea green), Percavirus, Rhadinovirus (light green) and a yellow star for the virus from this study. The heatmap illustrates the amino acid relative identity against this study novel virus across the different proteins used in the phylogeny. The colors correspond to the identity score, where dark purple indicates the lowest relative identity (0.4) and bright yellow indicates the highest relative identity (1.0). Node support is indicated by diamonds: black diamonds represent high support (SH-aLRT > 75 and UFBoot > 75), gray diamonds represent support superior to 75 in SH-aLRT only, and white diamonds in UFBoot only. The scale bar indicates the number of amino acid substitutions per site.

#### Adenoviridae

A total of 360 contigs were identified as belonging to the family *Adenoviridae*. These assemblies were also identified in *D. rotundus* under BioProjects PRJEB28138 and PRJEB34487. Contigs lengths varied from 103 bp to 4,079 bp, and all contigs were successfully mapped against *Mastadenovirus desmodi Vampire bat adenovirus* (BK066905), showing high nucleotide identity (mean: 93.54%, min: 75.4%, max: 100%). Contigs containing near complete DNA polymerase, hexon and penton genes from libraries ERR2756788, ERR2756803, and ERR3569494 were extracted to further characterize their genetic diversity. Phylogenetic analysis based on the polymerase gene revealed that the novel viruses form a well-supported sister lineage (SH-aLRT = 78.3, UFBoot = 95) to *Mastadenovirus desmodi* (BK066905) with a highly supported monophyletic clade (SH-aLRT = 100, UFBoot = 100) **(Figure 9A)**. While the polymerase gene of the novel viruses is highly conserved and shares high identity with previously reported viruses (93.6–99.3% nucleotide and 97.1–98% amino acid identity), the major capsid proteins present a distinct pattern. The hexon gene shares 79.4–79.8% of nucleotide and 86.3–86.3% of amino acid identity, and the penton gene shares 89.8% of nucleotide and 93.8% amino acid identity. Notably, the hexon N-terminal domain showed a higher concentration of mutation when compared to other regions **(Figure 9B)**.

**Figure 9.**
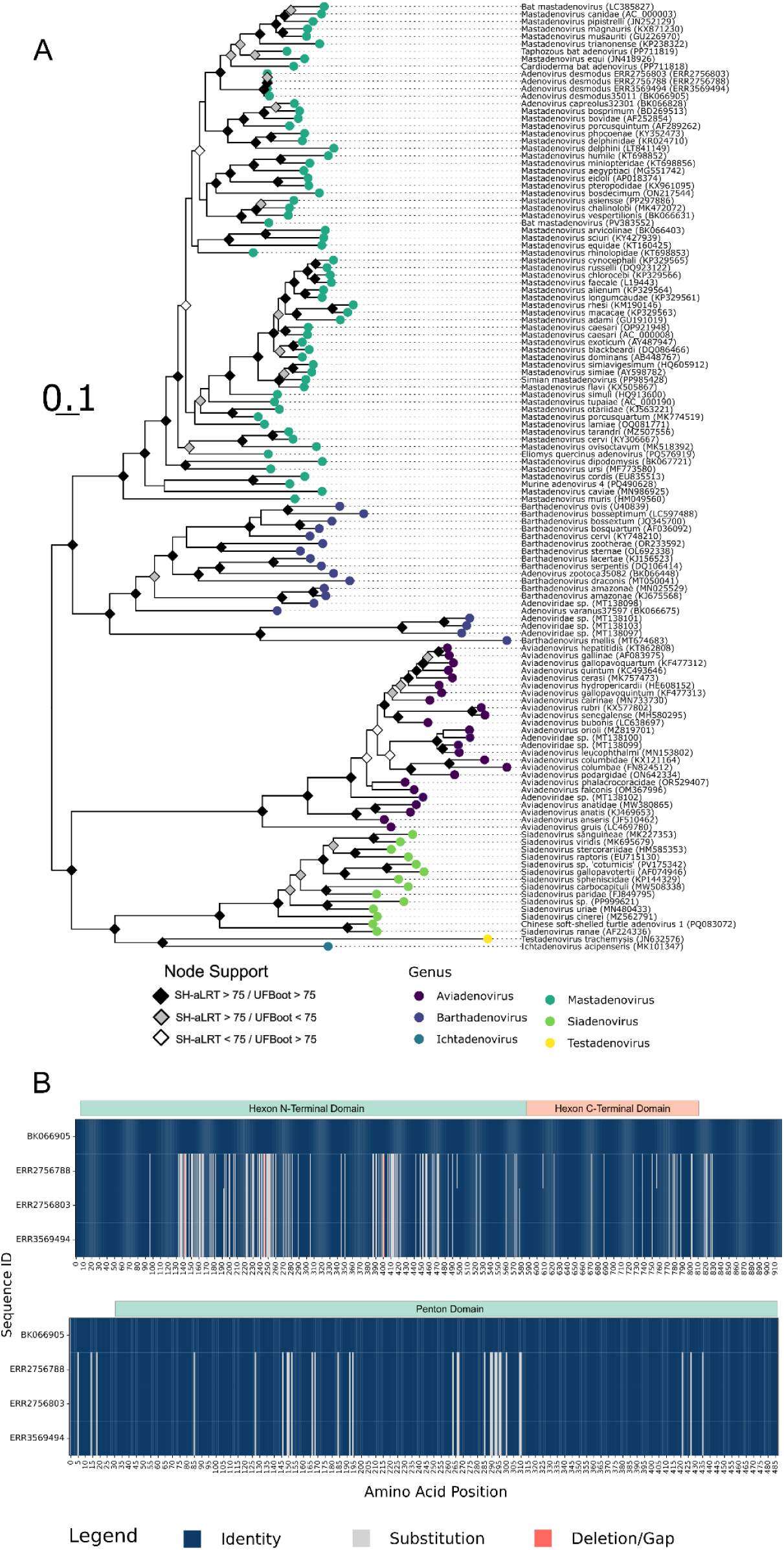
Phylogenetic and genetic characterization of a novel *Mastadenovirus* from *Desmodus rotundus*. (A) Maximum likelihood phylogenetic tree of the Adenoviridae family, based on the Polymerase gene, illustrating the phylogenetic placement of the novel viruses found in *Desmodus rotundus* (shown in teal). The tips of the tree are color-coded to represent different viral genera: *Mastadenovirus* (teal), *Aviadenovirus* (dark blue), *Batadenovirus* (purple), *Ichtadenovirus* (light blue), *Siadenovirus* (green), and *Testadenovirus* (yellow). Phylogeny was inferred with a 738 amino acids alignment under the model LG+I+G. Node support is indicated by diamonds: black diamonds represent high support (SH-aLRT > 75 and UFBoot > 75), gray diamonds represent support superior to 75 in SH-aLRT only, and white diamonds in UFBoot only. The scale bar indicates the number of amino acid substitutions per site. The scale bar indicates the number of amino acid substitutions per site. (B) Amino acid alignment map of the Hexon and Penton protein sequences from the novel viruses and the *Mastadenovirus desmodi* (BK066905) as reference. The plot visualizes positions of identity (dark blue), substitution (light blue), and deletion/gaps (pink). A notably higher concentration of mutations is observed in the N-terminal domain of the Hexon protein compared to other protein regions.

## Discussion

HTS is a powerful resource for expanding the virosphere ^1,13,50,51^. However, conventional discovery strategies frequently fail to capture the total viral diversity within a sample ^7^. Recent protein prediction or profiling approaches to detect the RdRp protein in generated HTS data demonstrate a vast diversity of viruses yet to be characterized ^52,53^. Nevertheless, a common bottleneck in viral discovery remains in the integration and scalability of bioinformatics. The existence of reference VDM discovery tools is key to comprehending this obscure viral diversity, however, they can require complex pre-processing steps, such as *de novo* assembly and the identification of contigs without any similarity to known biological sequences ^7,12,13^. To address these bottlenecks, we developed DeepVir, a Snakemake-based workflow ^17^ designed to automate and democratize the initial stages of viral discovery within HTS data. DeepVir provides a reproducible and scalable pipeline that transitions from raw HTS data to VDM identification, allowing researchers to prioritize biological interpretation of their findings rather than pipeline development and management.

To demonstrate the efficacy of DeepVir in a highly diverse yet under-sampled ecosystem, we analyzed the public SRA to investigate hidden viruses in American bats. This group of animals is known for its high diversity ^54,55^, and a well-documented role as reservoirs for viruses with zoonotic potential ^18,22,23,56–59^. However, regional sampling within America remains sparse ^60^. While PCR-based methods are traditionally favored due to their low cost and targeted nature ^61–65^, their reliance on conserved genomic structures makes them less effective at detecting divergent pathogens. Given viruses’ short generation time and high mutation rates, common diagnostics tools may fail during novel strain outbreaks ^66–68^. Recent systematic reviews evidenced biases of most PCR applications or specific agent tests in the process of viral discovery ^69^. Among the 26 viral families identified in bats in America, 83.2% of the 3,500 records point to three viral families: *Rhabdoviridae* (n = 2.361), *Coronaviridae* (n = 361), and *Paramyxoviridae* (n = 192) ^70^. This discrepancy highlights how negative results can be an artifact of testing methods, as well as of the study population, season, or sample type ^71^.

The robust nature of the DeepVir pipeline is evidenced by the identification of 179 distinct viral groups in the 203 paired-end SRA datasets of american bats, which were successfully characterized by phylogenetics. This significantly expanded the known viral diversity, with novel viral sequences belonging to well established lineages in RNA viruses (families *Orthomyxoviridae, Picornaviridae, Alphaflexiviridae, Astroviridae*, and *Coronaviridae*), reverse transcribing viruses (*Retroviridae*), and DNA viruses (families *Adenoviridae, Herpesviridae,* and *Papillomaviridae*), as well as 170 novel lineages. This success characterization stems from integration standard local alignment and homology-based search tools ^72–75^ for general virus identification, complemented by palmDB-centered RdRp identification (via *palm_annot*) ^76^. Despite the successful identification of VDM members, determining their ecological niches and putative hosts remains a major hurdle, as exemplified by the clusters 6 and 12. As novel viral lineages are continuously uncovered at a petabase scale ^7,13–15^, future efforts on developing models to elucidate the ecological roles and host interactions of these newly discovered entities are required.

Furthemore, among the identified viruses, only the family *Papillomaviridae*, *Adenoviridae*, along with the subfamily *Spumaretrovirinae* were reported in the original works, which explored and evaluated the zoonotic potential of novel viruses in the common vampire bat from Peru ^77,78^. Whereas remaining viral agents were not reported due the goal of exploring the transcriptome and evolution within bats in the original works ^79–82^. The large-scale application of DeepVir also enabled the characterization of known viral families of high zoonotic relevance, overcoming previous limitations caused by fragmented assemblies. Specifically for *Spumaretrovirinae*, overcoming these limitations and assembling the genomic puzzle was only possible through human-driven curation of the pipeline’s outputs, which allowed for the identification and linkage of homologous contigs spanning distinct genomic regions across multiple sequencing libraries. This approach allowed for the discovery of the first putative exogenous *Spumavirus* in the Americas ^83^. Furthermore, the pipeline’s sensitivity was highlighted by the detection of a near-complete Influenza A H7N9 genome in the cosmopolitan bat *Carollia perspicillata*. These findings underscore unknown transmission routes between avian and mammalian reservoirs and the discovery of novel *Picornaviridae* lineages that parallel Asian bat diversity, revealing a significant geographic gap in pathogen surveillance.

Beyond immediate zoonotic threats, the identified virosphere sheds light on complex host-virus evolutionary dynamics and environmental dissemination. The discovery of a novel *Gammaherpesvirinae* lineage and *Adenoviridae* contigs exhibiting localized capsid mutations suggests long-term codivergence and ongoing viral evasion of bat immune responses. Interestingly, the robust assembly of plant-associated *Alphaflexiviridae* genomes (e.g., *Lolavirus*) in frugivorous bats points to an unexplored mechanical or environmental transmission route, raising questions about the role of wildlife in the dissemination of agricultural pathogens.

The present work provides a pipeline to aid viral discovery in HTS data and further expands viral diversity in American bats, revealing viral agents hidden in previously published sequencing data ^79,80,82,84–86^. Nevertheless, to assess spillover potential within the virosphere of American fauna, comprehensive knowledge of viral diversity and robust detection tools are essential. Identifying the specific host species and their geographic distributions is pivotal for prioritizing surveillance efforts. Integrating HTS data into the monitoring of unknown pathogens, alongside the development of specific diagnostic tools and easy-to-use bioinformatic pipelines, represents a promising future approach for targeted pathogen surveillance ^87–89^. Mapping viral diversity in wildlife establishes the necessary baseline to identify past or future spillover events that have thus far gone undetected.

## Acknowledgments

The authors acknowledge the Bioinformatics Core Facility of the Brazilian National Cancer Institute (INCA) for their support and PhD Nicole Scherer for all the assistance during the project development. The authors acknowledge the ISO 9001-certified IRD i-Trop HPC (South Green Platform) at IRD Montpellier for providing HPC resources that have contributed to the research results reported in this paper. URLs: https://bioinfo.ird.fr and http://www.southgreen.fr. The authors thank PhD Martine Peeters for her valuable input and thoughtful review of the manuscript.

## Data Accessibility

As this study exclusively utilized publicly available, open-access HTS data obtained from the SRA (https://www.ncbi.nlm.nih.gov/sra), it did not involve the direct handling or sampling of animals by the authors. The DeepVir pipeline developed and used for the present work can be found in the public repository, https://github.com/matheus-cosentino/DeepVir. The scripts and generated files of SSN, network and VDM Taxonomic references can be found at Zenodo repository https://zenodo.org/records/18681598 (DOI: 10.5281/zenodo.18681598).

## Author contributions

Conceptualisation: MACC, MD, AFAS. Data curation: MACC. Formal analysis: MACC. Software: MACC, NFN, MD. Investigation: MACC, MD, AFAS, AA, NFN, MS. Resources: AA, MP, MS. Funding acquisition: MD, AFAS, AA, MACC. Supervision: MD, AA, AFAS. Project administration: MD, AA, MACC. Writing – Original Draft Preparation: MACC. Writing – Review and editing: MD, AFAS, AA, MF, NFN, MS.

## Supporting Information Captions

**Supporting Information Appendix S1 - Bioinformatic methods to contextualize novel viral agents identified by the DeepVir pipeline**

**Supporting Information Appendix S2 - Supplementary images to contextualize datasets and novel viruses identified by DeepVir.**

**Supplementary Table S1. - Summary of SRA Libraries.** Contains detailed information (S1.1), summary per species (S1.2), and summary per diet (S1.3) of the transcriptomic samples used in the study.

**Supplementary Table S2. - Metadata for phylogenetic datasets.** Includes accession numbers, organism names, and other metadata for reference sequences used to characterize Orthomyxoviridae, Picornaviridae, Alphaflexiviridae, Spumaretrovirinae, Orthoherpesviridae, Papillomaviridae, and Adenoviridae contigs.

**Supplementary Table S3. - Overall viral diversity and computational analysis outputs.** Contains summaries of identified viruses, Diamond BLASTx contig parameters, Palm_annot and Palmid outputs, MMseqs2 generated clusters, and Sequence Similarity Network (SSN) neighbor and cluster compositions.

**Supplementary Table S4. - Diamond BLASTx summary.** Detailed outputs for the identification of protein sequences within the contigs of Orthomyxoviridae, Picornaviridae, Alphaflexiviridae, Papillomaviridae, and Herpesviridae.

**Supplementary Table S5. - Protein identity matrices.** Matrices used for phylogenetic contextualization across different viral families and their respective proteins (e.g., 3Dpol, P1, Replicase, Capsid, L1, L2, E1, E2, DNA pol, DNA pack, Helicase, Envelope, Uracil).

**Supplementary Data S1. Phylogenetic datasets and sequence alignments.** Compressed archive (.zip) containing all sequence alignments and raw phylogenetic datasets generated and analyzed in this study.

