## Supporting Information Appendix for "DeepVir: A reproducible workflow for large-scale viral dark matter discovery"

Guia 1

**DiscoVir: A scalable snakemake pipeline unravels the American bat virosphere**

**Table of Contents:**

**Appendix S1: Bioinformatic methods to contextualize novel viral agents identified by the DiscoVir pipeline	3**

VDM Contextualization	3

Back-Mapping and Annotation	5

Phylogenetic Analysis	5

Influenza A Subtyping	6

**Appendix S2: Supplementary images to contextualize datasets and novel viruses identified by DiscoVir.	7**

**References	22**

### Appendix S1: Bioinformatic methods to contextualize novel viral agents identified by the DiscoVir pipeline

#### VDM Contextualization

After the DiscoVir preliminary results, to prevent the misidentification of the putative novel RNA VDM, contigs containing RdRp palm cores underwent a manual cross-validation step. First, original nucleotide sequences were searched against the nr NCBI nucleotide database v.5 (<https://ftp.ncbi.nlm.nih.gov/blast/db/v5/>) using BLASTn v.2.16.0 ^1^ to remove misidentified host transcripts. Subsequently, structural validation of putative ORFs encoding RdRp was performed using Phyre2 ^2^. Following the methodology described for RdRp-scan ^3^, contigs exhibiting high-confidence structural homology to known viral RdRps in the PDB were retained. Furthermore, sequences displaying structural hits with a confidence threshold below 90% to any entry were also retained as putative highly divergent viral dark matter. Conversely, sequences matching non-viral proteins with >90% confidence were discarded as false positives. All retained candidates were subsequently subjected to genomic annotation and evolutionary analysis.

The validated RdRp palm core domains were processed using the palmid workflow (<https://github.com/ababaian/palmid>) to identify closely related sequences within VDM, using the latest version within the PalmDB v.2023-04-26 ^4^ (available at <https://github.com/ababaian/palmdb>). Briefly, candidate sequences were subjected to homology searches against the PalmDB database using Diamond v.2.0.15 ^5^ in ultra-sensitive mode, with a maximum of 10 hits retained and low-complexity masking disabled. For each contig with significant hits, an individual multiple sequence alignment was generated using MAFFT v7.520 ^6^. Phylogenetic relationships were subsequently inferred using FastTreev.2.2. ^7^ to generate maximum-likelihood trees under the JTT+CAT model ^8^ and Branch support inferred by 1,000 replicates of Shimodaira–Hasegawa test ^7^.

All RdRp-containing contigs identified and validated were contextualized through a comparative structural approach using a Sequence Similarity Network (SSN) visualization. Initially, sequence redundancy was collapsed using MMseqs2 ^9^ (easy-cluster) with a 30% identity threshold and 80% coverage (-c 0.8 --cov-mode 1). The resulting clusters were then aligned using MAFFT v7.520 ^6^ and converted into Hidden Markov Model (HMM) profiles by the hh-suite HHmake v.3.3.0 ^10^. Subsequently, a high-sensitivity MMseqs2 all-against-all search (-s 7.5) was performed comparing all sequences against all HMM profiles. To estimate evolutionary divergence while accounting for alignment length and sequence variability, a normalized distance matrix was calculated following ^11^. This distance was defined as the negative natural logarithm of the ratio between the MMseqs2 profile-vs-sequence bit-score and the maximum bit-score achievable by the profile (**Formule**).

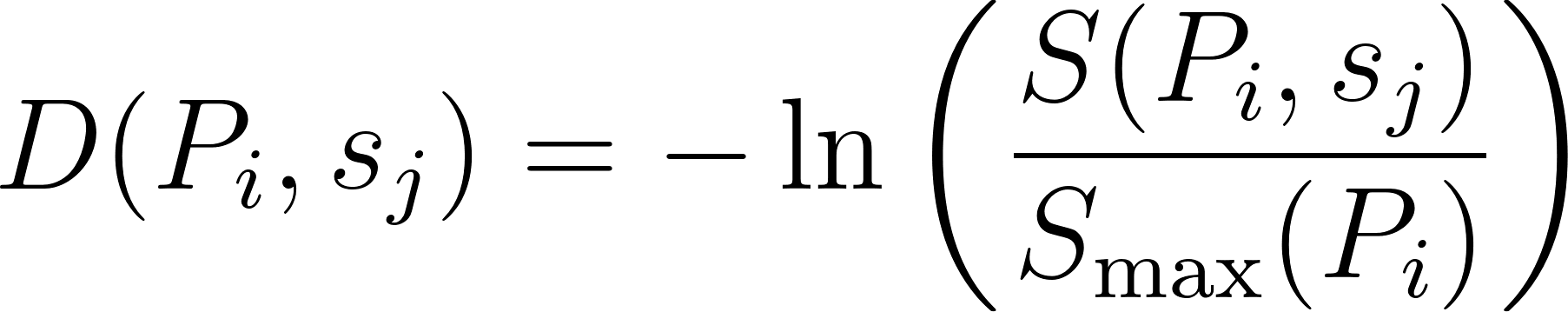
<https://www.codecogs.com/eqnedit.php?latex=D(P_i%2C%20s_j)%20%3D%20-%5Cln%20%5Cleft(%20%5Cfrac%7BS(P_i%2C%20s_j)%7D%7BS_%7B%5Cmax%7D(P_i)%7D%20%5Cright)#0>

##### Formule - Normalized distance matrix calculation based on bit-scores to quantify the evolutionary distance between highly divergent viral sequences. Sequence divergence (*D*) was estimated by calculating the negative natural logarithm of the ratio between the observed alignment score (*S(Pi,sj)*) to the maximum self-score of the query sequence (*Smax(Pi)*).

The distance matrix was imported into Python igraph library v.1.0.1 ^12^ to construct an undirected, weighted graph. The pairwise connections were defined as edge lists, and individual RdRp sequences as nodes for SSN visualization. Unsupervised community detection was performed using the Louvain algorithm ^13^, with network topology and node coordinates computed using the Distributed Recursive Graph Layout algorithm ^14^. To validate the biological relevance of the derived clusters and assess the taxonomic novelty of our dataset, reference RdRp palmcore sequences and metadata were retrieved from the Serratus project database ^15^. Reference nodes were annotated at the order and family levels. The taxonomic composition of each Louvain cluster was quantified by calculating the number and percentage of established taxa mapped within the resulting clusters.

#### Back-Mapping and Annotation

To confirm viral presence within each library, sequencing data were mapped to *de novo* assembled contigs following the ViralUnity workflow v.1.0.5 (available at <https://github.com/filiperomero2/ViralUnity>). Briefly, raw reads were filtered by quality (Q30) and length (50 bp) using Trimmomatic v.0.39 ^16^. Filtered reads were mapped using Minimap2 ^17^ and were sorted using SAMtools v.1.11 ^18^. Variant calling and consensus genome inference was performed using BEDtools v1.12 ^19^. When more than one sample type per individual was available, the estimated viral load was expressed as reads per million (RPM) and compared between sample types using the Mann-Whitney U test ^20^.

#### Phylogenetic Analysis

Validated contigs passed by Prokka workflow v.1.14.5 ^21^ and Geneious Prime software v.2024.0.5 TransferTools (<https://www.geneious.com>) to predict putative genes within assembled data. Predicted genes of viral contigs identified by taxonomic assignment with Diamond were evolutionarily contextualized. Local databases of viral proteins of interest were filtered by Diamond BLASTx and were aligned against reference sequences ([Supplementary Table S2](https://docs.google.com/spreadsheets/u/0/d/1QNdZMU-hRaBsqJf6XugYrsHUuvHuBJr0cU6BwUbbceI/edit)). Phylogenetic informative regions were translated using Aliview v.1.28 ^22^ and aligned by amino acid information using MAFFT v7.520 ^6^. Alignment regions rich in gaps were removed using a heuristic selection based on similarity statistics in trimal v.1.4.rev15 (-automated1) ^23^. The final alignment was used to infer a maximum-likelihood tree by IQ-Tree v.2.2.5 ^24^ under the best-fitting amino acid substitution model selected by ModelFinder ^25^. Branch support was inferred by 10,000 replicates of UltrafastBootstrap (UFboot) and 10,000 replicates of approximate Likelihood Ratio Test and Shimodaira–Hasegawa (SH-aLRT) ^26^. Phylogenetic trees were visualized and annotated using the ggtree v.3.10.1 R package ^27^. Identity matrices of amino acid alignments were calculated using bio3d v.2.4-5 R package ^28^.

#### Influenza A Subtyping

Identified Influenza A sequences recovered through data mining were subjected to subtyping by performing multiple sequence alignments and inferring maximum likelihood phylogenies for Hemaglutinin (H) and Neuraminidase (NA) genes using similar methods as described in the section 4.4. Reference sequences were obtained at the Bacterial And Viral Bioinformatics Resource Center ^29^. To this end, individual nucleotide sequences were translated into amino acids through AliView v.1.30 ^22^ and were aligned with MAFFT v7.520 ^6^. After that, the amino acid alignment was transformed into a nucleotide alignment though backTranslate.py, an in-house script available at <https://github.com/matheus-cosentino/EvoTools>. Sequences were trimmed and used to infer the maximum-likelihood nucleotide genotype phylogeny, according to ^30^.

### Appendix S2: Supplementary images to contextualize datasets and novel viruses identified by DiscoVir.

##### Supplementary Figure S1 - Summary of chiropteran BioSamples used for present study. (A) Total SRA by Species, (B) Samples by Family, and (C) Samples by Tissue. The graphs are color-coded according to the diet of the respective species: Frugivory (dark purple), Hematophagous (dark red), Insectivore (teal), Nectarivore (dark green), Omnivorous (light green), and Piscivory (yellow).

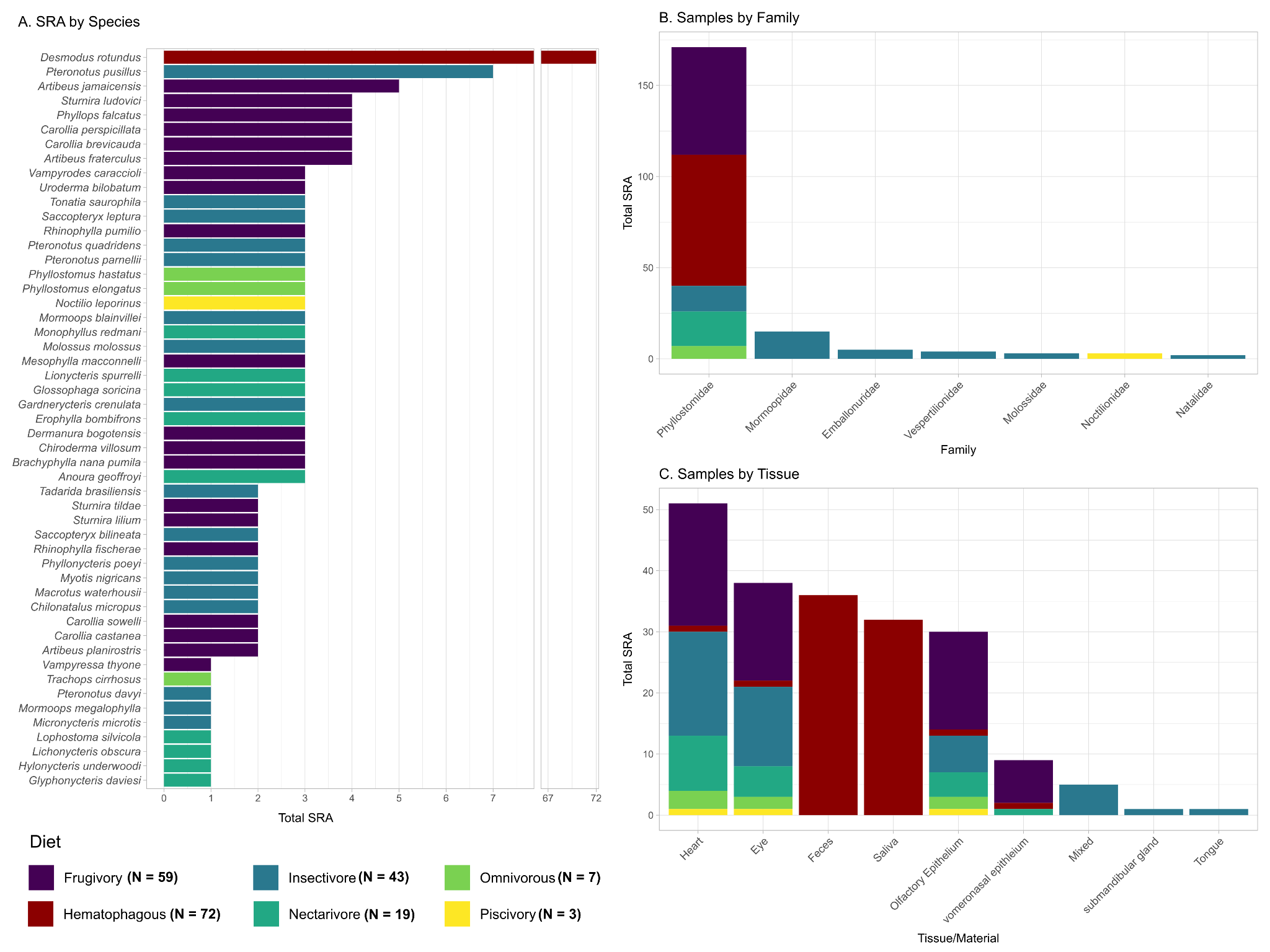

##### Supplementary Figure S2 - Overall PalmID phylogenetic tree of identified viral contigs. Maximum likelihood phylogenetic tree illustrating the placement of the assembled contigs (highlighted in red). Nodes are colored by SH-aLRT support above 70, represented by the white nodes. The scale bar indicates the number of amino acid substitutions per site.

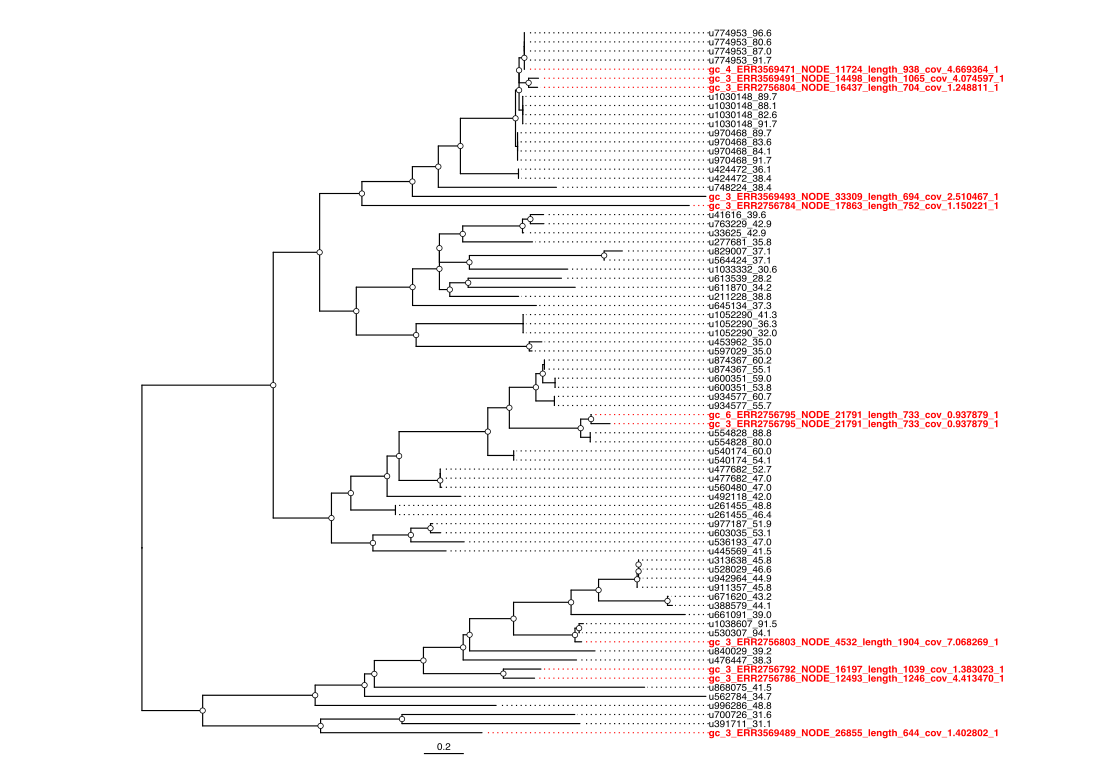

##### Supplementary Figure S3 - Individual PalmID phylogenetic subtrees of identified viral contigs. Detailed phylogenetic subtrees showing the evolutionary relationships of specific assembled contigs (highlighted in red). Nodes are colored by SH-aLRT support above 70, represented by the white nodes. The scale bar indicates the number of amino acid substitutions per site.

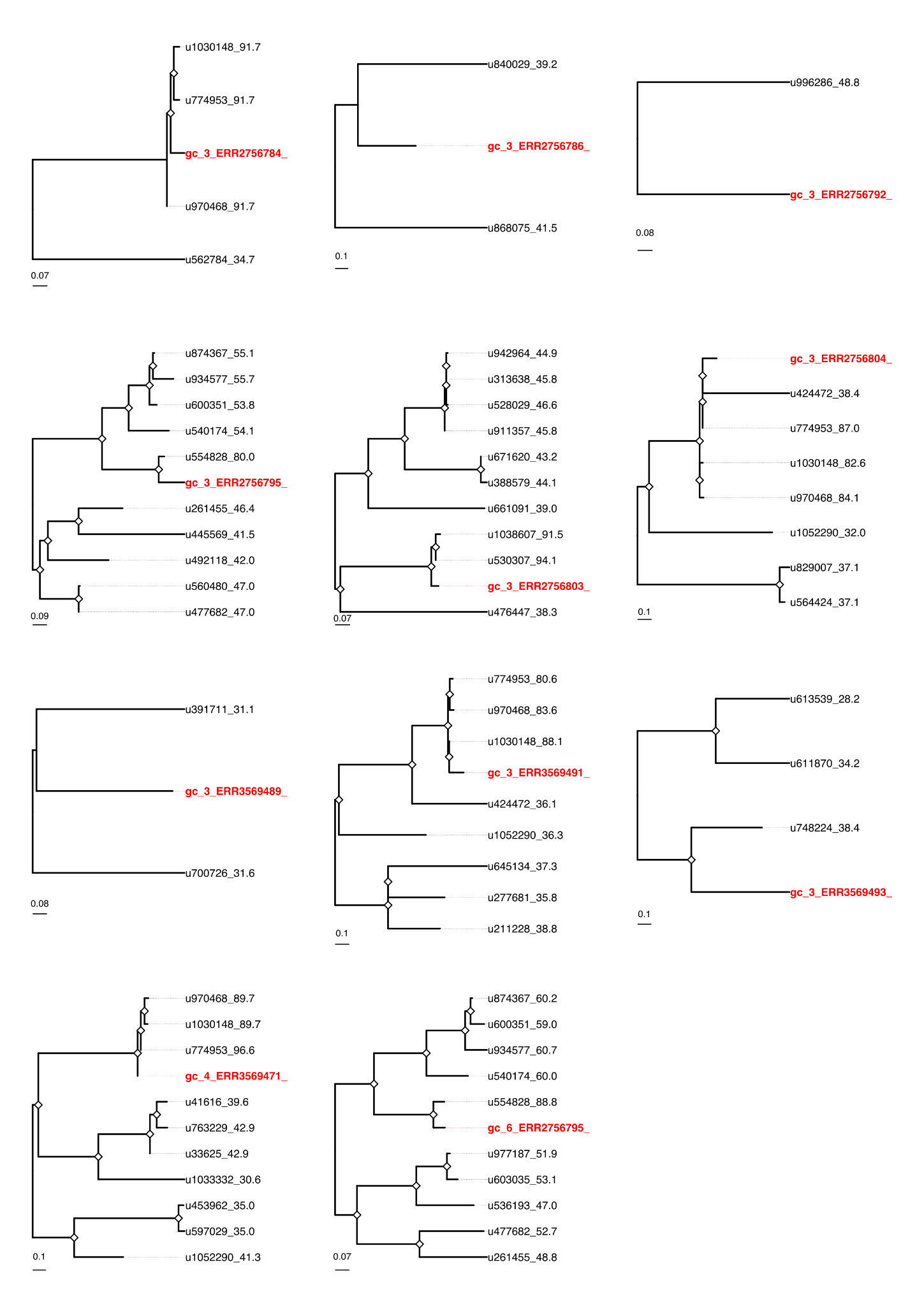

##### Supplementary Figure S4 - Sequence Similarity Network (SSN) individual clusters visualized by Taxonomic Order composition. Structural homology-based SSN visualization of VDM sequences partitioned into Louvain clusters (0, 2 to 11). Nodes represent individual RdRp sequences. The nodes are color-coded according to their annotated taxonomic Order to evaluate the dominant taxa within each cluster and infer putative host associations and ecological niches. Sequences generated in this study are highlighted in red, allowing for the visual contextualization of these novel viruses alongside known viral orders.

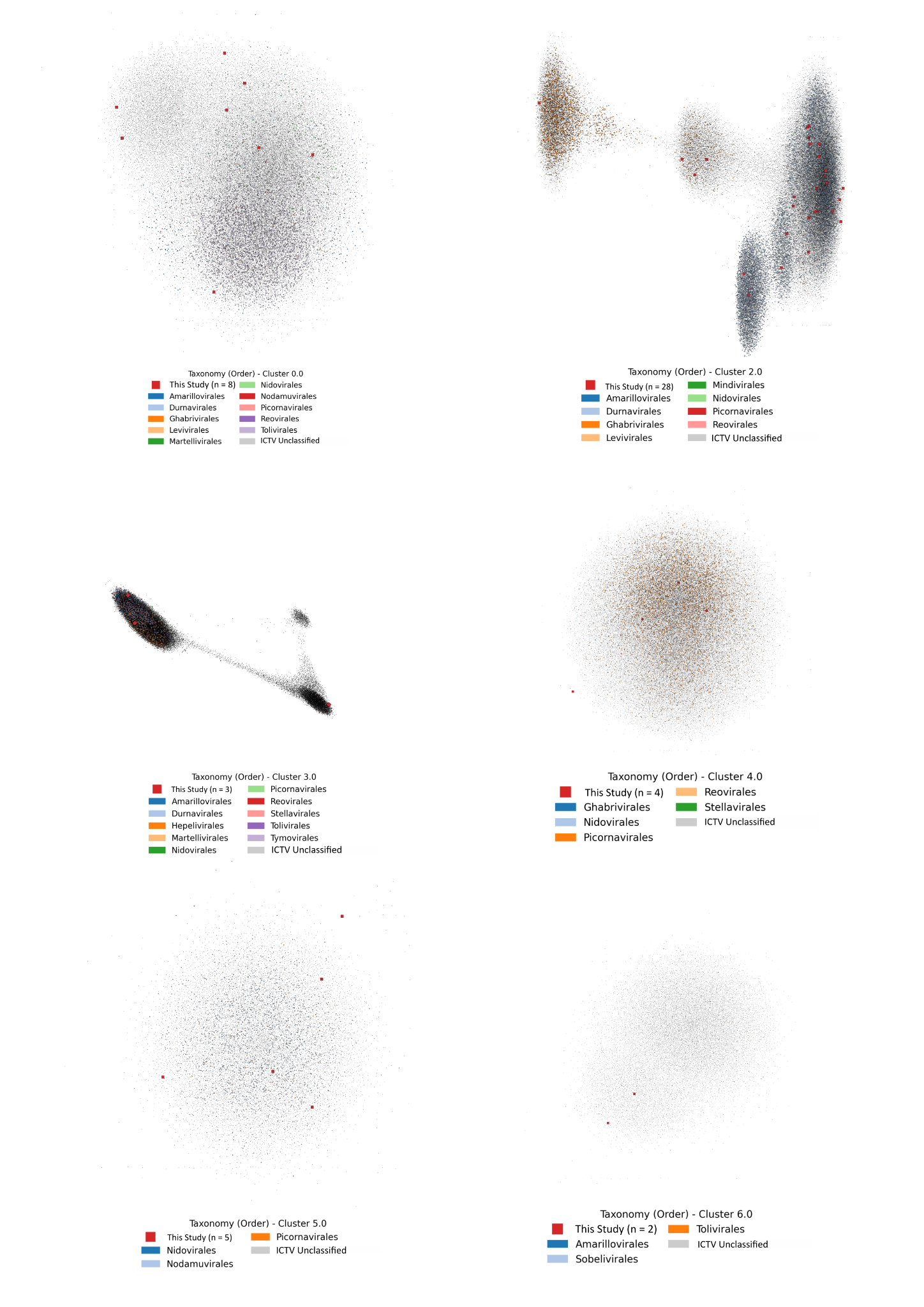

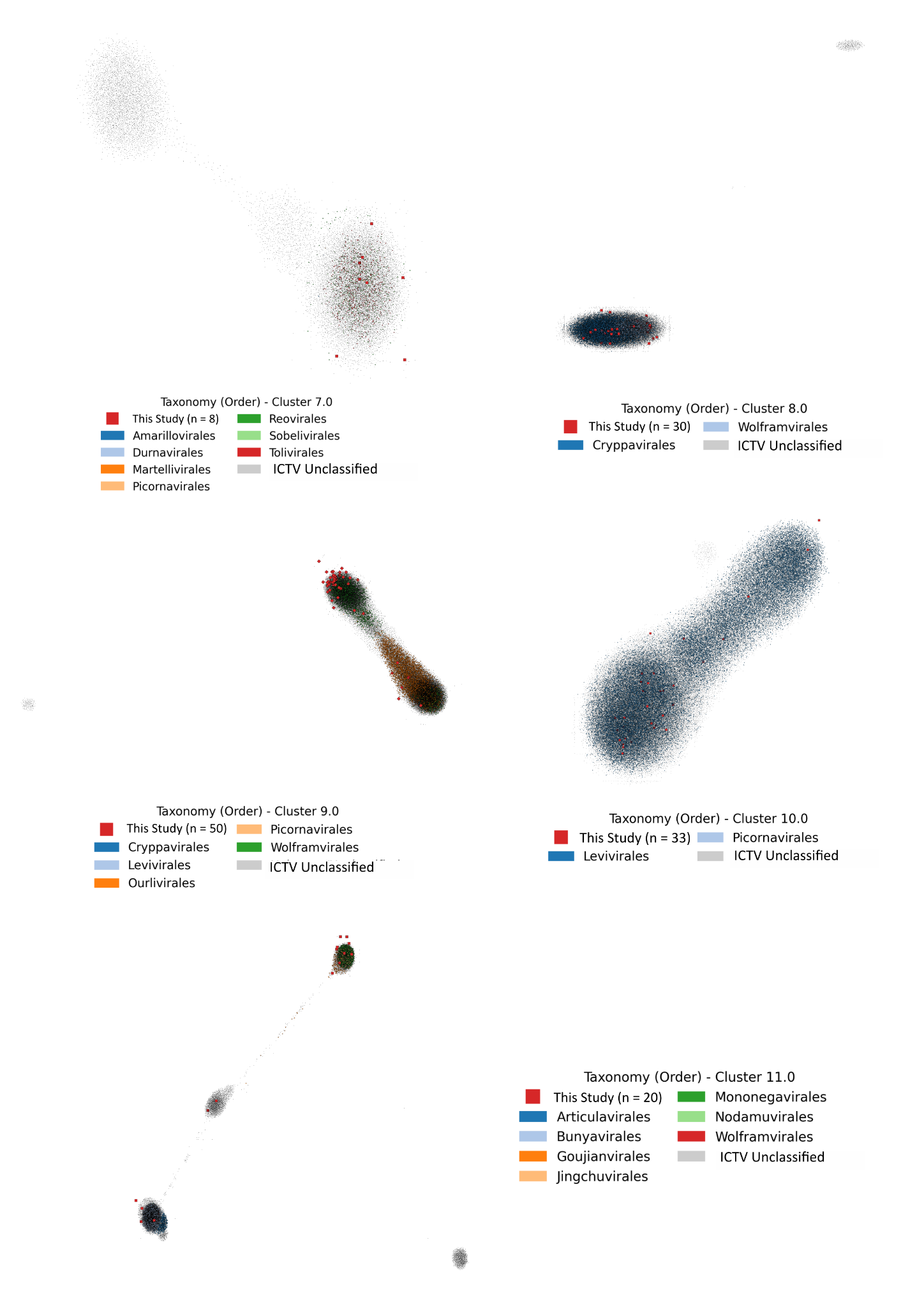

##### Supplementary Figure S5 - Sequence Similarity Network (SSN) individual clusters visualized by Taxonomic Family composition. Structural homology-based SSN visualization of VDM sequences partitioned into Louvain clusters (0, 2 to 11). Nodes represent individual RdRp sequences and are color-coded according to their annotated taxonomic family. Sequences generated in this study are highlighted in red, allowing for the visual contextualization of these novel viruses alongside known viral orders.

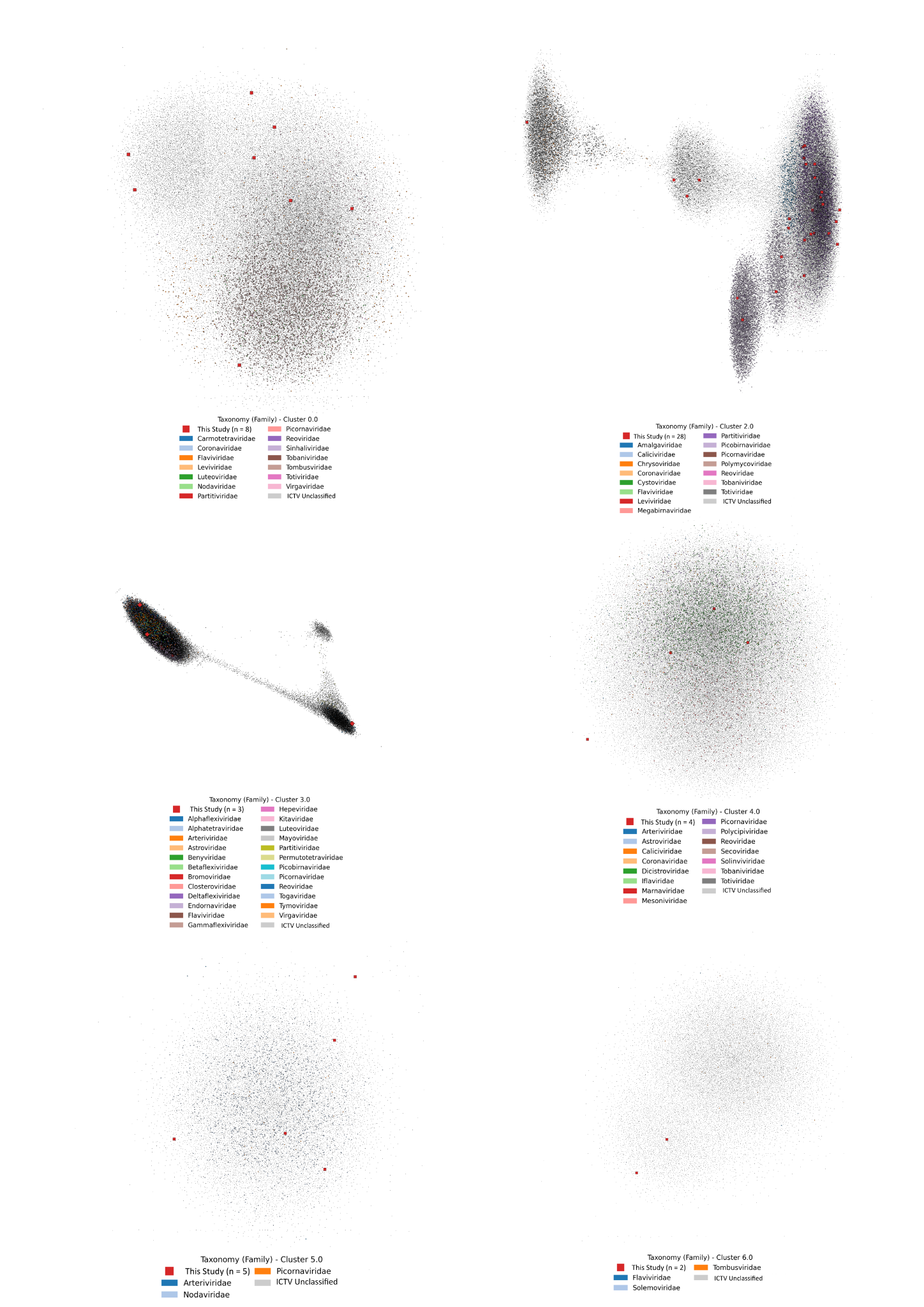

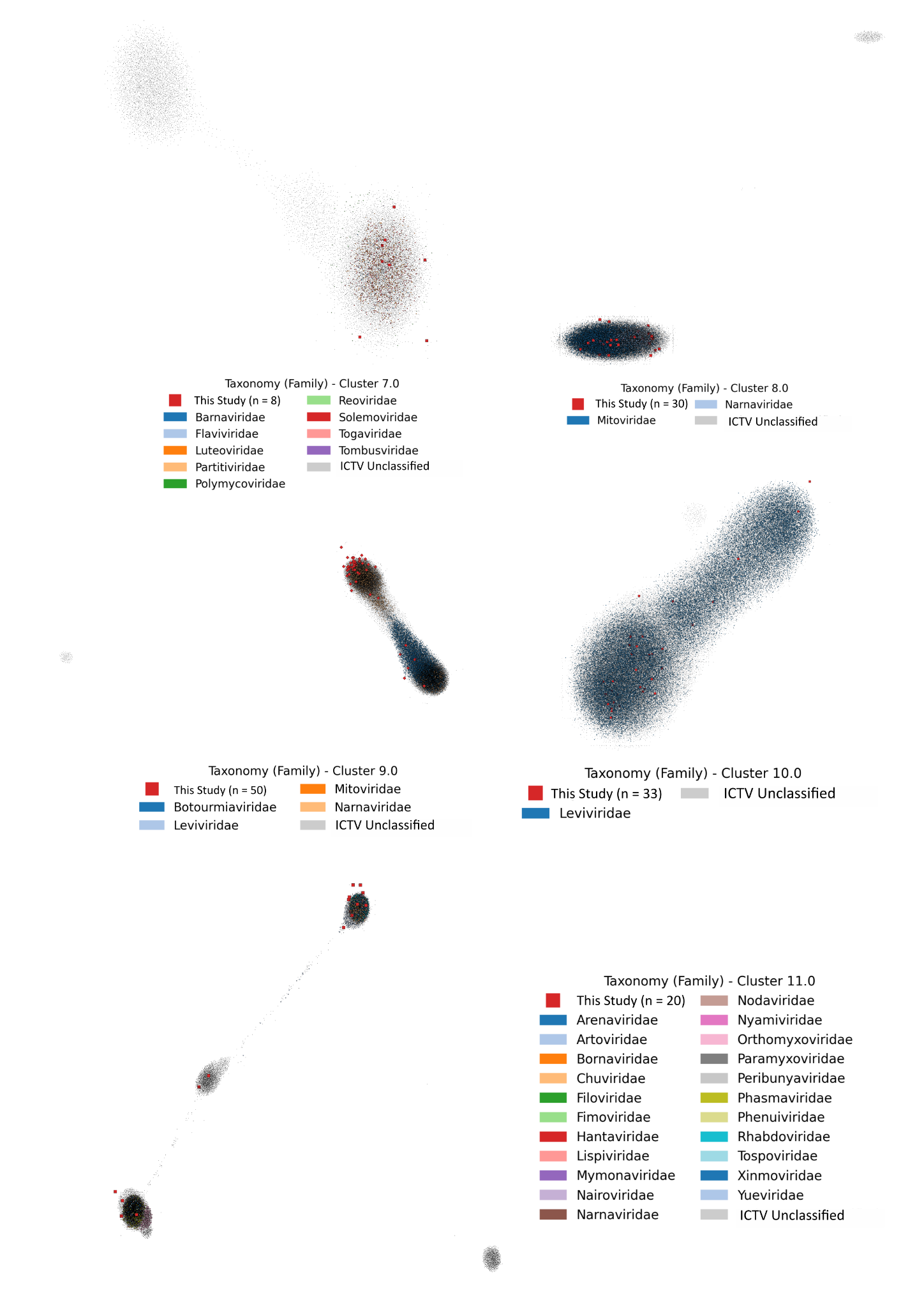

##### Supplementary Figure S6 - Maximum likelihood phylogenetic tree of PB1 sequences from Orthomyxoviridae. The tree illustrates the phylogenetic placement of the newly identified sequences within the Orthomyxoviridae family. Phylogeny was inferred from an alignment of 14 sequences and 544 amino acid sites under the Q.PFAM+F+I+G4 model. The tips are color-coded by Genus: Alphainfluenzavirus (purple), Betainfluenzavirus (dark blue), Gammainfluenzavirus (teal), Isavirus (dark green), Quaranjavirus (light green), and Thogotovirus (yellow). Sequences of this study are highlighted with a purple star. Node support is indicated by diamonds: black diamonds represent high support (SH-aLRT > 75 and UFBoot > 75), gray diamonds represent support superior to 75 in SH-aLRT only, and white diamonds in UFBoot only. The scale bar indicates the number of amino acid substitutions per site.

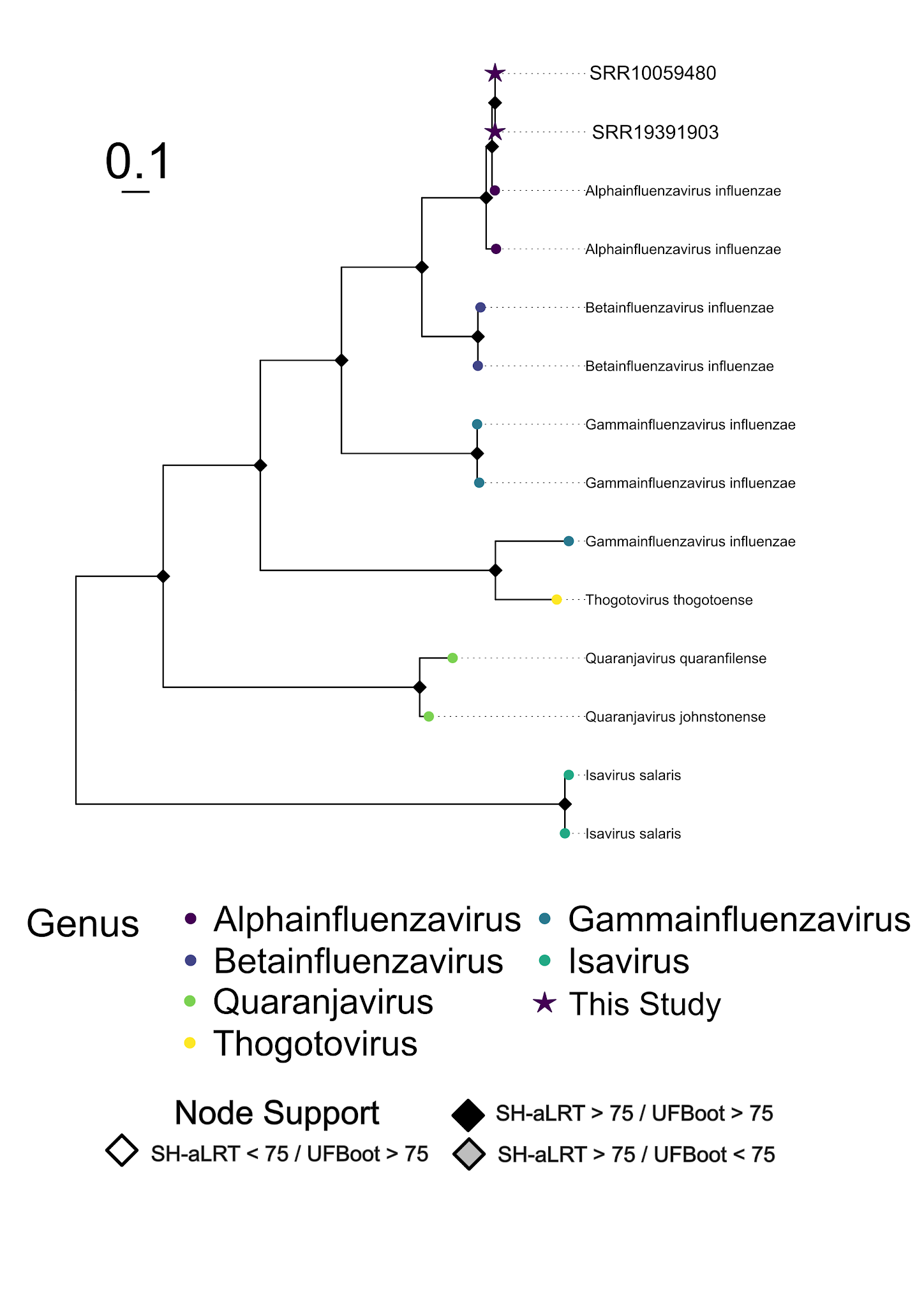

##### Supplementary Figure S7 - Maximum likelihood phylogenetic trees for Influenza A subtyping. (A) Phylogeny based on the Hemagglutinin (HA) gene, inferred from 810 sequences and 1,758 nucleotide sites under the GTR+F+I+G4 model. (B) Phylogeny based on the Neuraminidase (NA) gene, inferred from 804 sequences and 981 nucleotide sites under the GTR+F+I+G4 model. Sequences of this study are highlighted with a grey star. The scale bar indicates the number of amino acid substitutions per site. Node support is indicated by diamonds: black diamonds represent high support (SH-aLRT > 75 and UFBoot > 75), gray diamonds represent support superior to 75 in SH-aLRT only, and white diamonds in UFBoot only. The scale bar indicates the number of amino acid substitutions per site.

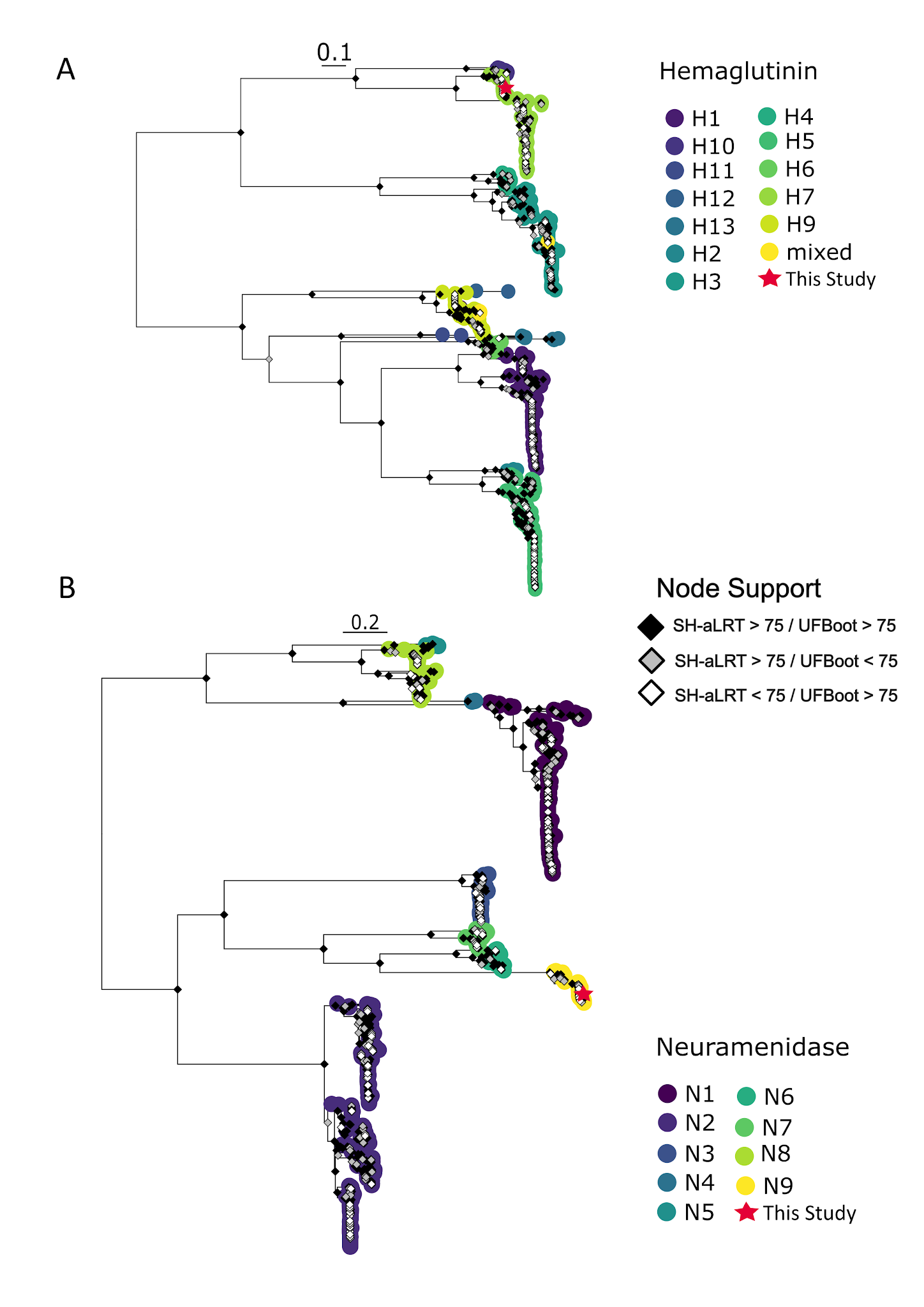

##### Supplementary Figure S8 - Schematic representation of Coronaviridae and Astroviridae fragments identified in public databases. (A) Genomic organization of the Coronaviridae contigs mapped against the reference, Mimon bat coronavirus isolate PREDICT/PDF-3316 (MZ293744). The color bar indicates the percentage of nucleotide identity, ranging from lower identity in dark purple (~87%) to higher identity in yellow (100%). (B) Genomic organization of the Astroviridae contigs mapped against the reference, Jingmen bat astrovirus 1 (OQ802697). The color bar indicates the shared amino acid identity, ranging from lower identity in dark purple (~35%) to higher identity in yellow (~60%).

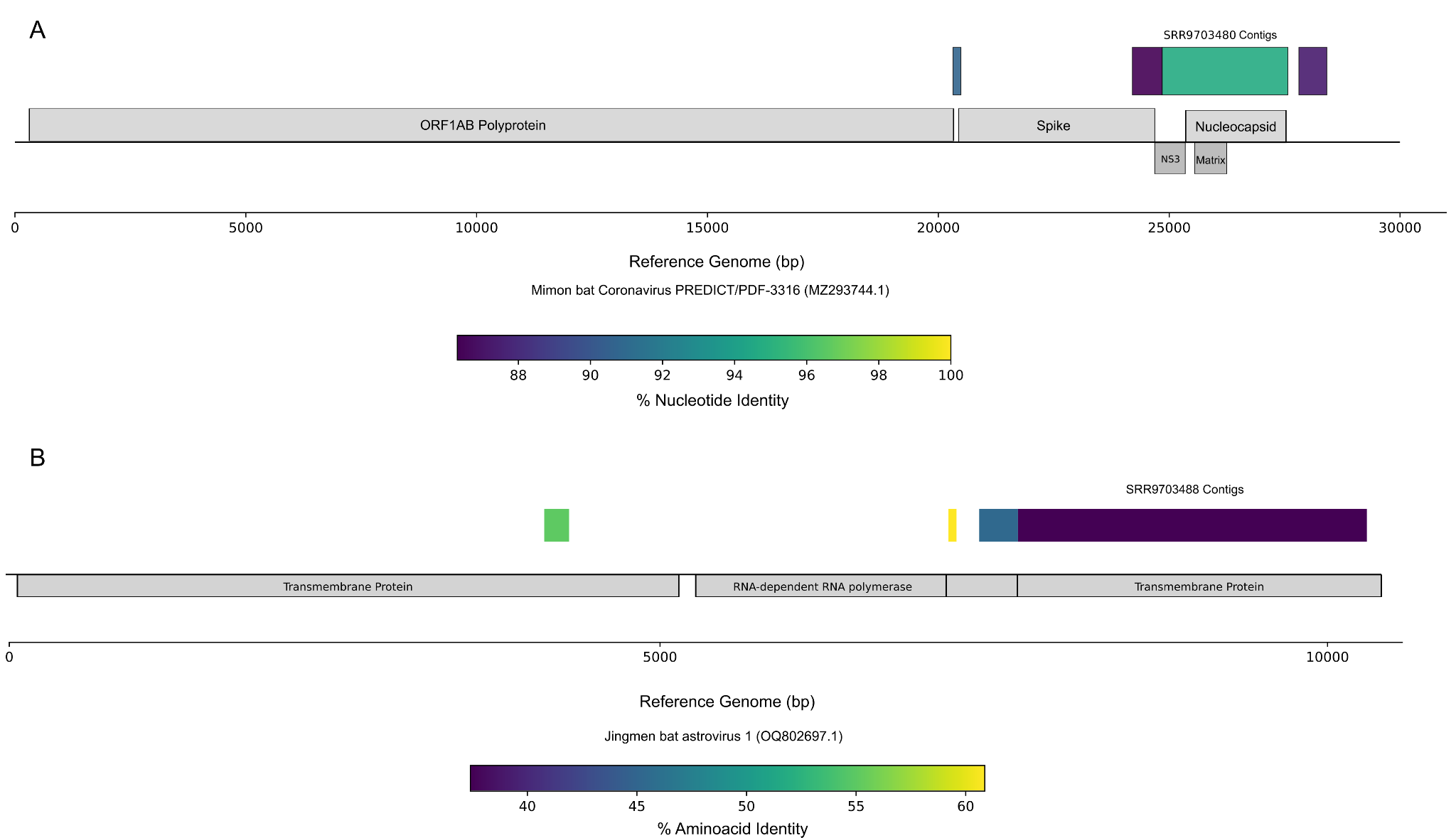

##### Supplementary Figure S9 - Maximum likelihood phylogenetic tree of Papillomaviridae L1 gene found within American bats. Maximum likelihood phylogenetic tree of the Papillomaviridae family, based on the L1 genes, illustrating the phylogenetic placement of the novel viruses found. Viruses found in the present work are highlighted in dark blue. Phylogeny was inferred with an alignment of 611 sequences and 200 amino acids under the model Q.insect+I+G4. The tips of the tree are color-coded to represent different hosts: Chiroptera (dark blue), Carnivora (purple), Others (blue), Primates (teal), Rodentia (dark green) and Sirenia (light green). Node support is indicated by diamonds: black diamonds represent high support (SH-aLRT > 75 and UFBoot > 75), gray diamonds represent support superior to 75 in SH-aLRT only, and white diamonds in UFBoot only. The scale bar indicates the number of amino acid substitutions per site.

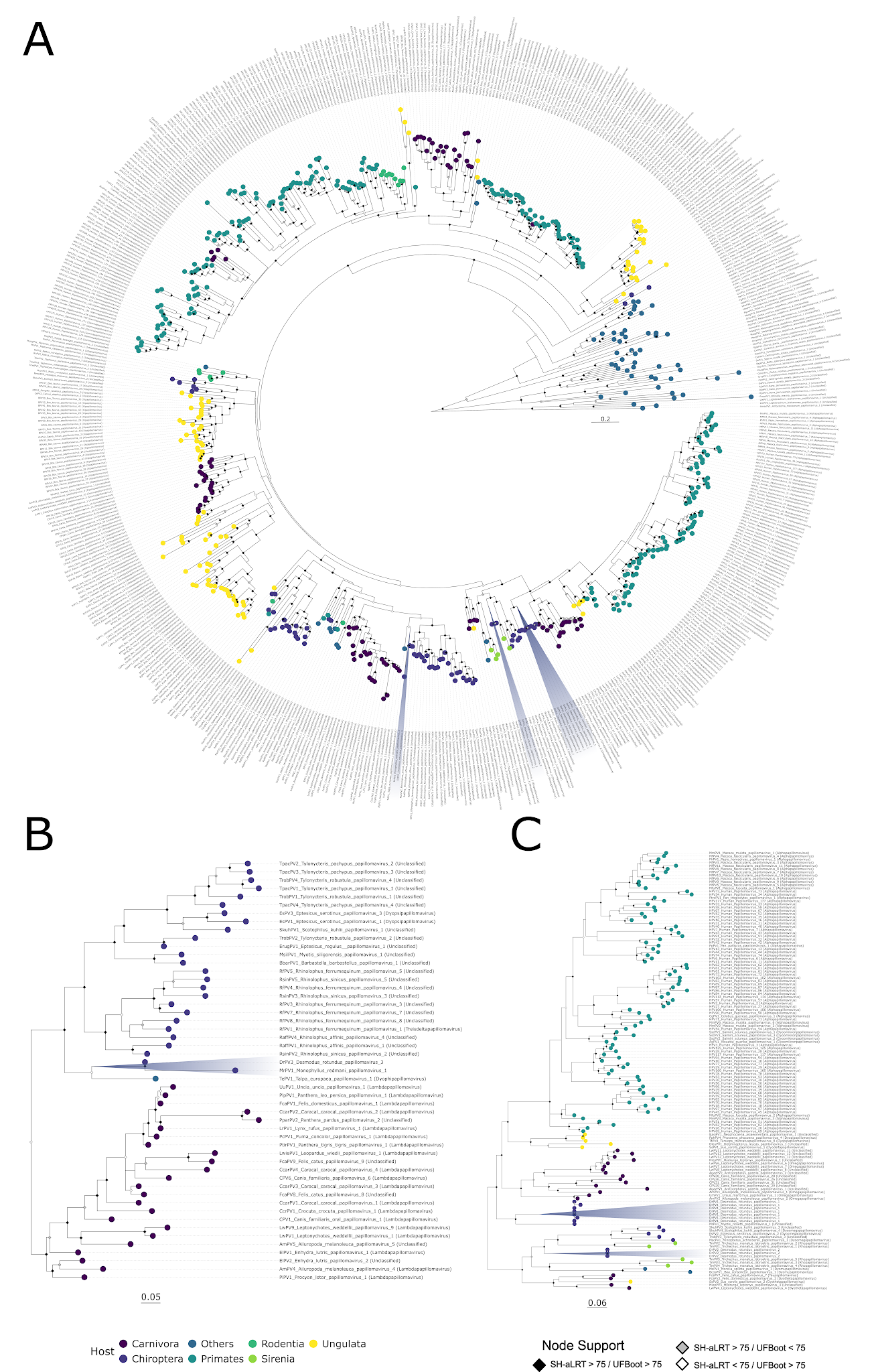

### Supporting Information References

1.	Camacho, C. *et al.* BLAST+: architecture and applications. *BMC Bioinformatics* 10, 421 (2009).

2.	Kelley, L. A., Mezulis, S., Yates, C. M., Wass, M. N. & Sternberg, M. J. E. The Phyre2 web portal for protein modeling, prediction and analysis. *Nat. Protoc.* 10, 845–858 (2015).

3.	Charon, J., Buchmann, J. P., Sadiq, S. & Holmes, E. C. RdRp-scan: A bioinformatic resource to identify and annotate divergent RNA viruses in metagenomic sequence data. *Virus Evol* 8, veac082 (2022).

4.	Edgar, R. C. *et al.* Petabase-scale sequence alignment catalyses viral discovery. *Nature* 602, 142–147 (2022).

5.	Buchfink, B., Reuter, K. & Drost, H.-G. Sensitive protein alignments at tree-of-life scale using DIAMOND. *Nat. Methods* 18, 366–368 (2021).

6.	Katoh, K. & Standley, D. M. MAFFT multiple sequence alignment software version 7: improvements in performance and usability. *Mol. Biol. Evol.* 30, 772–780 (2013).

7.	Price, M. N., Dehal, P. S. & Arkin, A. P. FastTree 2--approximately maximum-likelihood trees for large alignments. *PLoS One* 5, e9490 (2010).

20.	Website. https://www.jstor.org/stable/2236101?seq=2.
